# The dimorphic diaspore model *Aethionema arabicum* (Brassicaceae): Distinct molecular and morphological control of responses to parental and germination temperatures

**DOI:** 10.1101/2023.12.14.571707

**Authors:** Jake O. Chandler, Per K.I. Wilhelmsson, Noe Fernandez-Pozo, Kai Graeber, Waheed Arshad, Marta Pérez, Tina Steinbrecher, Kristian K. Ullrich, Thu-Phuong Nguyen, Zsuzsanna Mérai, Klaus Mummenhoff, Günter Theißen, Miroslav Strnad, Ortrun Mittelsten Scheid, M. Eric Schranz, Ivan Petřík, Danuše Tarkowská, Ondřej Novák, Stefan A. Rensing, Gerhard Leubner-Metzger

## Abstract

Plants in habitats with unpredictable conditions are often characterized by diversifying their bet-hedging strategies that ensure fitness over a wider range of variable environmental factors. A striking example is the diaspore (seed and fruit) heteromorphism that evolved to maximize species survival in *Aethionema arabicum* (Brassicaceae) in which external and endogenous triggers allow the production of two distinct diaspores on the same plant. Using this dimorphic diaspore model, we identified contrasting molecular, biophysical, and ecophysiological mechanisms in the germination responses to different temperatures of the mucilaginous seeds (M^+^ seed morphs), the dispersed indehiscent fruits (IND fruit morphs), and the bare non-mucilaginous M^−^ seeds obtained by pericarp (fruit coat) removal from IND fruits. Large-scale comparative transcriptome and hormone analyses of M^+^ seeds, IND fruits, and M^−^ seeds provided comprehensive datasets for their distinct thermal responses. Morph-specific differences in co-expressed gene modules in seeds, as well as seed and pericarp hormone contents identified a role of the IND pericarp in imposing coat dormancy by generating hypoxia affecting ABA sensitivity. This involved expression of morph-specific transcription factors, hypoxia response and cell wall-remodeling genes, as well as altered abscisic acid (ABA) metabolism, transport, and signaling. Parental temperature affected ABA contents and ABA-related gene expression and altered IND pericarp biomechanical properties. Elucidating the molecular framework underlying the diaspore heteromorphism can provide insight into developmental responses to globally changing temperatures.

**IN A NUTSHELL:** *Background:* Heteromorphic diaspores (fruits and seeds) are an adaptive bet-hedging strategy to ensure survival in spatiotemporally variable environments. The stone cress *Aethionema arabicum*, an annual plant native to semi-arid habitats in Anatolia (Turkey), one of the world’s hotspots of biodiversity. It is a close relative of Arabidopsis, rapeseed, cabbage and other *Brassica* crops, but in contrast to these *Ae. arabicum* disperses two distinct diaspores from the same plant. These dimorphic diaspores are the mucilaginous seeds (dispersed by pod shatter) and indehiscent fruits (dispersed by abscission). The wing-like pericarp (fruit coat) of the single-seeded indehiscent fruit allows wind dispersal over large distances. The amounts and ratios of the dimorphic diaspores are variable and depend on the environmental conditions. The dimorphic diaspores differ in morphology, dormancy and germination properties and thereby make *Ae. arabicum* an excellent model for the comparative investigation of the underpinning molecular mechanisms.

*Question:* We asked how temperature during fruit and seed formation and during seed germination affect dormancy release and germination speed, and how the morphology, hormonal regulation, and the expression of genes differ between the dimorphic diaspores.

*Findings:* Large-scale comparative transcriptome and hormone analyses of the mucilaginous seeds and the indehiscent fruits, as well as the seeds artificially extracted from indehiscent fruits by pericarp (fruit coat) removal, provided comprehensive datasets for their distinct thermal responses. Material obtained from plants grown at different temperatures during reproduction was imbibed at different temperatures for germination. This altered the abscisic acid (ABA) metabolism and the pericarp biomechanical properties. Diaspore-specific differences in response to distinct imbibition temperatures identified distinct gene expression patterns in seeds, distinct seed and pericarp hormone contents, and a role of the pericarp in generating hypoxia inside the fruit and imposing coat dormancy. This revealed distinct combinations of specific transcription factors, hypoxia responses and cell wall-remodeling genes, as well as altered signaling pathway genes.

*Next steps:* Our large-scale comparative transcriptome datasets are easily and publicly accessible via the *Aethionema arabicum* web portal (https://plantcode.cup.uni-freiburg.de/aetar_db/index.php). We plan to expand this by future work on seedlings derived from the dimorphic diaspores, by comparing different *Ae. arabicum* genotypes, and by studying responses to specific stresses. Understanding the molecular basis of this fascinating example of developmental diversity and plasticity and its regulation by temperature is expected to add insight how plants respond to changing environmental conditions.

## Introduction

Fruits and seeds as propagation and dispersal units (diaspores) have evolved an outstanding diversity and specialization of morphological, physiological, and biomechanical features during angiosperm evolution. Coordination of diaspore maturation as well as of diaspore germination timing with environmental conditions is essential for the critical phase of establishing the next generation of plants (Finch-Savage and Leubner-Metzger, 2006; Donohue et al., 2010). This is especially critical in annual species that must establish germination and plant growth in a given season or persist as diaspores in the seedbank for germination in a later season (Finch-Savage and Footitt, 2017). Seed dormancy, i.e., innate block(s) to the completion of germination of an intact viable diaspore under favorable conditions, is the key regulatory mechanism involved in this timing. Temperature during plant reproduction (parental growth temperature) and temperature sensing by the dispersed diaspore provide input determining dormancy depth, germination timing, and adaptation to climatic change (Walck et al., 2011; Fernandez-Pascual et al., 2019; Batlla et al., 2022; Iwasaki et al., 2022; Zhang et al., 2022).

Most species with dry fruits, including *Arabidopsis thaliana* (Brassicaceae), produce seed diaspores released by dehiscence, spontaneous opening at preformed structures from mature fruits (Mühlhausen et al., 2013). Other species have dry indehiscent fruits where one or more seeds remain encased by the pericarp (fruit coat). These indehiscent fruits are dispersed by abscission, exemplified by several Brassicaceae species (Lu et al., 2015b; Sperber et al., 2017; Mohammed et al., 2019). The pericarp of these indehiscent fruit diaspores may confer coat-imposed dormancy and delayed germination of the enclosed seeds. While most plants have evolved single types of diaspores that are optimized to the respective habitat, other plants employ a bet-hedging strategy by producing different types of diaspores on the same individual plant.

In these cases of diaspore heteromorphism, seeds and fruits differ in morphology, dormancy and germination properties, ecophysiology, and/or tolerance to biotic and abiotic stresses (Imbert, 2002; Baskin et al., 2014; Gianella et al., 2021). This diversity maximizes the persistence of a species in environments with variable and unpredictable conditions. Diaspore heteromorphism evolved independently in 26 angiosperm families and is common in the Asteraceae, Amaranthaceae, and Brassicaceae. Examples of seed dimorphism include the black and brown seed morphs of *Chenopodium album* and *Suaeda salsa* (Amaranthaceae), which differ in dormancy and responses to salinity (Baskin et al., 2014; Liu et al., 2018; Loades et al., 2023). The *Cakile* clade (Brassicaceae) produces fully indehiscent or segmented, partially indehiscent fruits (Hall et al., 2006). The dimorphic desert annual *Diptychocarpus strictus* (Brassicaceae) disperses short-lived winged, mucilaginous seeds and long-lived indehiscent siliques each containing about 11 seeds (Lu et al., 2015a). While the ecophysiology of these three dimorphic species is well described, the underpinning molecular mechanisms remain largely unknown.

As a model system to investigate the principles of diaspore dimorphism, we have chosen *Aethionema arabicum*, a small, diploid, annual, herbaceous species in the sister lineage of the core Brassicaceae, in which seed and fruit dimorphism was associated with a switch to an annual life history (Mohammadin et al., 2017). Genome and transcriptome information is available (Haudry et al., 2013; Nguyen et al., 2019; Wilhelmsson et al., 2019; Arshad et al., 2021; Fernandez-Pozo et al., 2021). *Aethionema arabicum* is adapted to arid and semiarid environments. Its life-history strategy appears to be a blend of bet-hedging and plasticity (Bhattacharya et al., 2019), and it exhibits true seed and fruit dimorphism with no intermediate morphs (Lenser et al., 2016). Two distinct fruit types are produced on the same fruiting inflorescence (infructescence): dehiscent (DEH) fruits with four to six mucilaginous (M^+^) seeds, and indehiscent (IND) fruits each containing a single non-mucilaginous (M^−^) seed. Upon maturity, DEH fruits shatter, releasing the M^+^ seeds, while the dry IND fruits are dispersed in their entirety by abscission. Dimorphic fruits and seeds differ in their transcriptomes throughout their development and in the mature dry state upon dispersal, and the dimorphic diaspores (M^+^ seeds and IND fruits) differ in their water uptake patterns and germination timing (Lenser et al., 2018; Arshad et al., 2019; Merai et al., 2019; Wilhelmsson et al., 2019; Nichols et al., 2020; Arshad et al., 2021). Together, these features qualify *Ae. arabicum* as a suitable model to investigate the molecular and genetic base of diaspore dimorphism.

Temperature is a main ambient factor affecting reproduction, dormancy and germination of plants (Walck et al., 2011; Fernandez-Pascual et al., 2019; Batlla et al., 2022; Iwasaki et al., 2022; Zhang et al., 2022) and temperature during reproductive growth is known to affect the ratio of IND/DEH fruit production of *Ae. arabicum* (Lenser et al., 2016). In our large-scale biology study, we provide a comprehensive comparative analysis of gene expression levels, hormonal status, biophysical and morphological properties underpinning the distinct *Ae. arabicum* dimorphic diaspore responses to ambient temperatures. By comparing M^+^ seeds dispersed from the fruits by dehiscence, indehiscent fruits containing M^−^ seeds dispersed by abscission, and bare M^−^ seeds obtained from IND fruits by manually removing the pericarp, we show that growth temperature during reproduction of the parent plant and a wide range of imbibition temperatures either promote or delay germination. We demonstrate how the pericarp of the IND fruit morph imposes coat dormancy.

## Results

### *Aethionema arabicum* reproductive plasticity and morph-specific responses to parental and imbibition temperatures

As described in the introduction, *Ae. arabicum* disperses two morphologically distinct diaspores (morphs), namely M^+^ seeds and IND fruits, that are produced at the same inflorescence (Figure 1A). The larger dehiscent fruits (DEH) release several M^+^ seeds upon maturation by dehiscence, whereas the smaller indehiscent fruits (IND) each containing a single M^−^ seed are dispersed by abscission (Figure 1A). Previous work (Lenser et al., 2016) showed that a 5°C increase in the ambient temperature during reproduction reduced the overall number of fruits and shifted the ratio between the two fruit types towards the DEH type. This parental temperature effect was confirmed here in a large-scale experiment with ca. 2000 plants at two parental temperature regimes during reproduction (20°C and 25°C; Figure 1B, Supplemental Figure S1A and Table S1). In earlier work (Lenser et al., 2016) we used 14°C as imbibition temperature to compare the germination and water uptake kinetics of the dimorphic diaspores from the 20°C parental temperature (PT) regime (20M^+^ seeds and 20IND fruits). This demonstrated that the germination of seeds enclosed in the 20IND fruits was much delayed compared to 20M^+^ seeds and bare 20M^−^ seeds obtained from the artificial separation of 20IND fruits by pericarp removal (Figure 1A). In the IND fruit, the pericarp makes up 74.4% of the morph’s mass but at maturity does not contain living cells (Arshad et al., 2019; Arshad et al., 2020; Arshad et al., 2021). The maximal germination percentage (G_max_) of the 20IND fruits was also much reduced compared to 20M^+^ and 20M^−^ seeds imbibed at 14°C (Lenser et al., 2016), indicating that the pericarp may impose coat dormancy.

**Figure 1.**
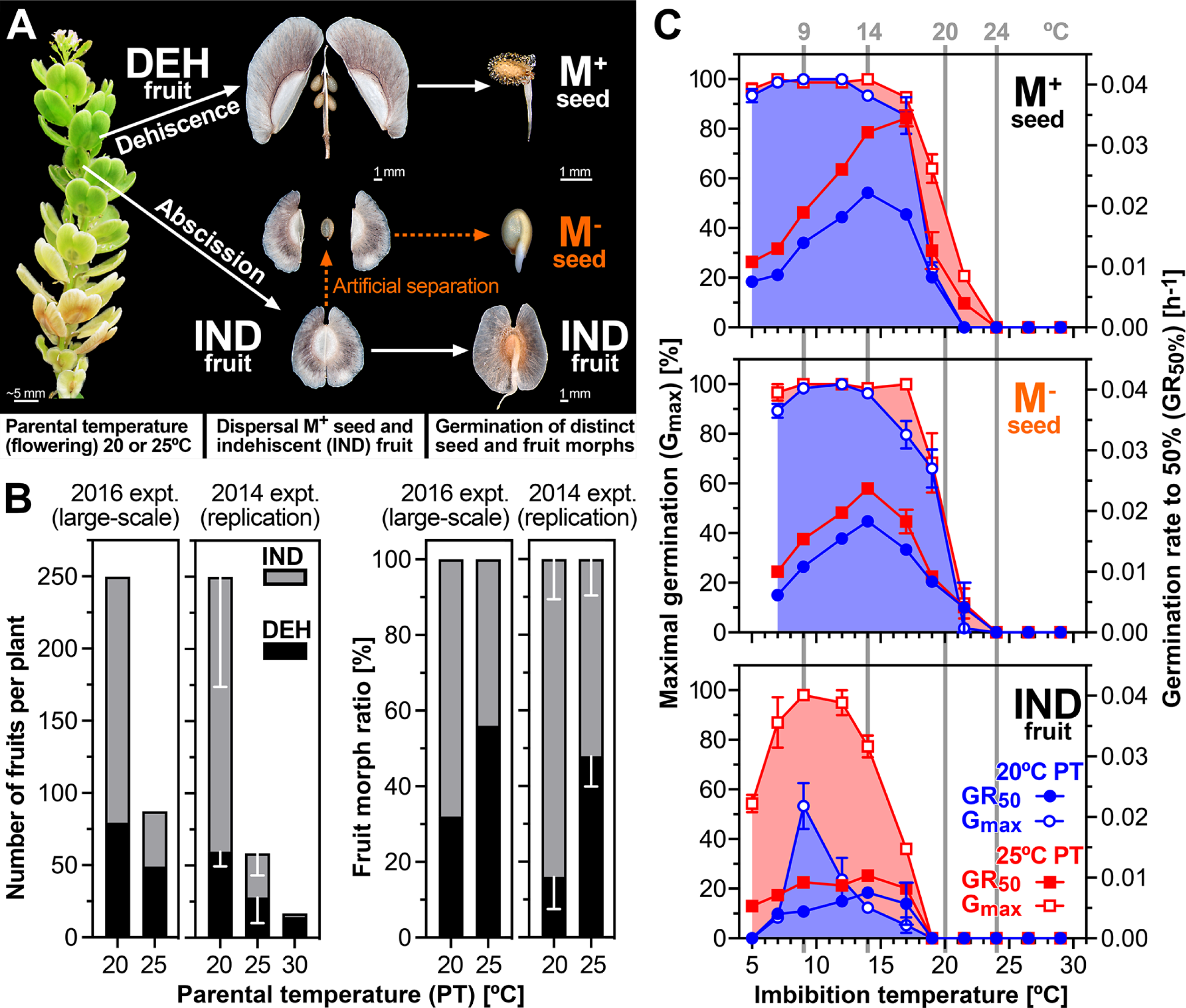
Dimorphic diaspore responses of *Aethionema arabicum* to ambient temperatures. A, Infructescence showing two morphologically distinct fruit types. Large, dehiscent (DEH) fruits contain four to six seed diaspores that produce mucilage (M^+^) upon imbibition. Small, indehiscent (IND) fruits contain a single non-mucilaginous (M^−^) seed each. For experiments with the bare M^−^ seed the pericarp was manually removed. B, The effect of parental temperatures (PT; ambient temperatures during reproduction) on the numbers and ratios of the fruit morphs in the 2016 harvest (large-scale) experiment (Supplemental Figure S1) and the 2014 harvest experiment (mean ± SD values of 3 replicates; total numbers of fruits were normalized to the large-scale experiment to aid comparison of the relative numbers for IND and DEH; the 20°C and 25°C 2014 harvest was used in the Lenser et al. (2016) publication). C, The effect of imbibition temperatures on the maximal germination percentages (G_max_) and the speed of germination expressed as germination rate (GR_50%_) of the dimorphic diaspores (M^+^ seeds, IND fruits), and for comparison of bare M^−^ seeds (extracted from IND fruits by pericarp removal). Sampling temperatures for molecular analyses are indicated. Mean ± SEM values of 3 replicate plates each with 20 seeds.

Here, we used the material generated in the large-scale experiment (Figure 1B) with two distinct parental temperatures, but otherwise identical growth conditions to address several aspects of the mechanisms underlying the diaspore dimorphism, especially the pericarp-imposed dormancy in a wide range of imbibition temperatures (Figure 1C). To investigate this pericarp effect more closer, we compared the germination-permissive temperature windows of freshly harvested mature M^+^ seeds with IND fruits as well as with bare M^−^ seeds (M^−^ obtained from IND by pericarp removal), obtained from plants grown at either 20°C (20M^+^, 20IND, 20M^−^) or 25°C (25M^+^, 25IND, 25M^−^) during reproduction. Seeds and fruits were imbibed at a range of temperatures between 5°C and 30°C, and their G_max_ as well as their germination rates (GR_50%_ = 1/T_50%_, a measure for the speed of germination with T_50%_ being the time required to reach 50% G_max_) were quantified (Figure 1C). IND fruits from plants matured at 20°C had a slower germination speed (lower germination rate, GR_50%_), a lower G_max_ at germination-permissive temperatures, and a narrower temperature range allowing near optimal germination, compared to M^+^ seeds from the same parents. For example, the 20IND fruits reached their highest G_max_ (ca. 50%) at 9°C but imbibed at even 2.5°C higher or lower, only reached 25% germination. On the other hand, the corresponding 20M^+^ seeds reached a G_max_ of above 85% from 5°C to 17.5°C. Pericarp removal demonstrated that 20M^−^ seeds had a similar optimum germination window as the 20M^+^ seeds, confirming the role of the IND pericarp in blocking germination (coat dormancy) at otherwise permissive temperatures for M^+^ and M^−^ seed germination.

M^+^ and M^−^ seeds from plants grown at the 20°C or 25°C parental temperature regimes had similar germination kinetics, although 25M^+^ and 25M^−^ seeds generally had a higher GR_50%_, and 25M^+^ seeds germinated slightly better at supra-optimal (warmer) temperatures (Figure 1C). Interestingly, the 25M^+^ seeds germinated much faster (t_50%_ 29 h) than 20M^+^ seeds when imbibed at 17°C (T_50%_ 54 h). Most different were 25IND fruits, which had a much higher G_max_ at germination-permissive temperatures than 20IND fruits (Figure 1C). 25IND fruits reached 87% to 98% germination at temperatures between 7°C and 12°C. Nonetheless, removal of the pericarp (IND *vs* M^−^) increased G_max_ and GR_50%_, especially at supra-optimal temperatures, for example, from 77% to 98% at 14°C and from 36% to 100% at 17°C. The pericarp therefore narrowed the permissive germination window by ca. 5°C at supra-optimal imbibition temperatures irrespective of the parental temperature. Thus, pericarp-imposed dormancy was still evident, although less extreme in 25IND fruits compared to 20IND fruits. At germination-permissive imbibition temperatures, the 25IND and 20IND pericarps therefore differed in their coat dormancy-imposing capabilities.

### Large-scale RNAseq and hormone quantification to identify morph-specific germination and dormancy mechanisms in *Ae. arabicum*

The contrasting pericarp-imposed dormancy and germination kinetics of M^+^, IND, and M^−^ showed that the two morphs integrate the signal of ambient imbibition temperature differently and suggest that one component is the ambient temperature during the reproduction of the parental plant (Figure 1). We hypothesized that the different germination responses to imbibition temperatures are mediated, at least in part, by transcriptional and hormonal changes during imbibition. We therefore collected M^+^, M^−^ and IND samples from plants grown at 20°C and 25°C parental temperature, respectively, during imbibition at four representative temperatures (9°C, 14°C, 20°C, and 24°C). In the sampling scheme (Supplemental Figure S1B), we considered physical (dry seed, 24 h imbibition) and physiological (T_1%_) times representing the progression of germination. 9°C is the most germination-permissive (G_max_) temperature for all morphs, still with a strong pericarp effect for 20IND (Figure 1C, Supplemental Figure S1C). At 14°C, M^+^ and M^−^ seed germination is permitted, while particularly 20IND fruit germination is inhibited more than at 9°C. This temperature was, therefore, chosen to examine the effect of the pericarp further. Imbibition at 20°C represents conditions when germination of all morphs is relatively inhibited, although a parental effect is evident, as 25M^+^ seeds germinate more readily than 20M^+^ seeds. At 24°C, germination of all diaspores is completely inhibited.

In the RNAseq analyses, counts of transcripts for 23594 genes of *Ae. arabicum* genome version 2.5 (Haudry et al., 2013) were obtained for each sample (Supplemental Data Set 1). To make the transcript abundance data easily and publicly accessible, we generated a gene expression atlas which is implemented in the *Ae. arabicum* genome database (DB) (Fernandez-Pozo et al., 2021) at https://plantcode.cup.uni-freiburg.de/aetar_db/index.php. This tool is based on EasyGDB, a system to develop genomics portals and gene expression atlases, which facilitates the maintenance and integration of new data and tools in future (Fernandez-Pozo and Bombarely, 2022). The *Ae. arabicum* gene expression atlas is very interactive and user-friendly, with tools to compare several genes simultaneously and multiple visualization methods to explore gene expression. It includes the transcriptome results of this work (135 datasets), of *Ae. arabicum* RNAseq work published earlier (Merai et al., 2019; Wilhelmsson et al., 2019; Arshad et al., 2021), and allows adding future transcriptome datasets. It also links to the newest version 3.1 of the *Ae. arabicum* genome annotation and sequence DB (Fernandez-Pozo et al., 2021) and allows linking to any improved future genome version. Further details and examples for the *Ae. arabicum* gene expression atlas, its analysis and visualization tools are presented in Supplemental Figure S2.

Principal component analysis (PCA) based on Log (normalized counts) from 22200 of 23594 genes after removing those with zero counts was used to observe general trends in the transcriptomes across the collected samples (Figure 2, Supplemental Figure S3). Prior to imbibition, there were differences in the dry seed transcriptomes, and although these samples cluster together in negative PC1 and PC2 coordinates (bottom left, Area A, Figure 2), a modest number of differentially expressed genes (DEGs) were identified between the samples (Supplemental Data Set 1). For example, there are 322 DEGs between the dry 20M^+^ seed and dry 25M^+^ seed transcriptomes, and 580 DEGs between dry 25M^+^ and dry 25M^−^ seed transcriptomes (Area A, Figure 2). Therefore, parental temperature and the pericarp (IND *vs* DEH fruit development) therefore affected the dry seed transcriptomes. A broad trend observed was that following increasing imbibition time: samples from seeds sown under generally germination-permissive conditions traveled positively along PC1 (to Areas C and D, Figure 2). Samples from seeds sown under germination-inhibiting conditions stay relatively closer to dry seeds, further supporting that PC1 represents ‘progress towards completing germination’ (Areas B, E and F, Figure 2). Reinforcing this, gene expression at an early imbibition timepoint of the 20M^+^ seeds is closer to the ‘dry seed’ state (Area B, Figure 2). The association between PC1 and final germination percentage is also evident in Supplemental Figure S3, which provides an extended PCA analysis.

**Figure 2.**
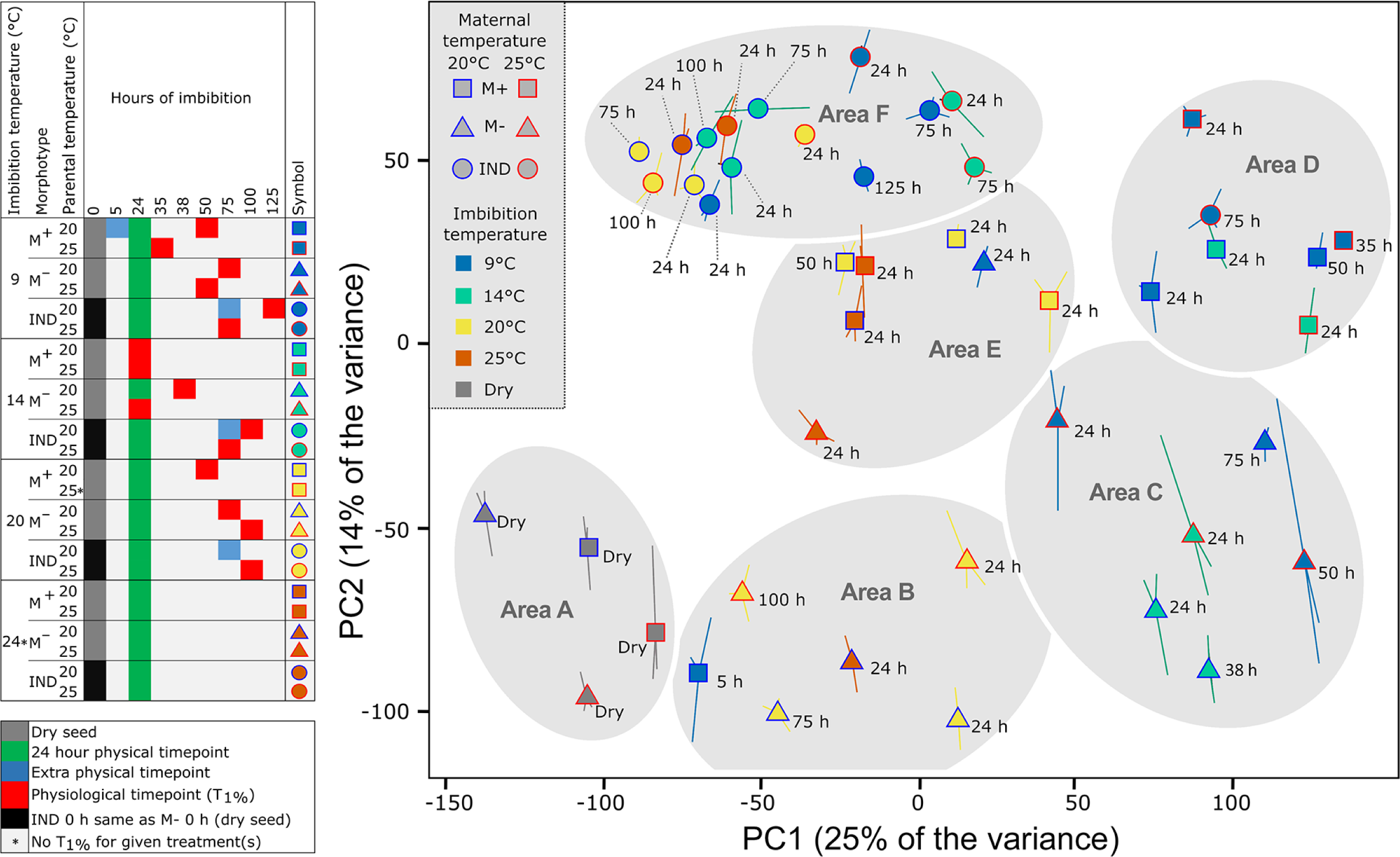
Principle components analysis (PCA) comparing the seed mRNA transcriptome data (RNA-sequencing analysis) of *Aethionema arabicum*. Mature M^+^ and M^−^ seeds, and IND fruits harvested from plants at two different parental temperatures during reproduction (20 and 25°C) were sampled in the dry state, and in the imbibed state at four different imbibition temperatures (9, 14, 20 and 24°C) and times indicated (e.g. 24 h); physiological time points (T_1%_) are also indicated. Indicated by asterisk, no germination occurred at 24°C imbibition temperature (precluding T_1%_ sampling) and 20M^+^ imbibed at 20°C was sampled only at 24h. PC1 and PC2 explain 25% and 14% of the variance; for PC3 and individual samples see Supplemental Figure S3. Large points indicate average coordinates from three replicates, with the location of each replicate relative to the average shown with a line (some lines are hidden by large point), time point label drop line differentiated by dotted line.

PC2 appears to generally separate IND and M^+^ from bare M^−^ seed, suggesting that PC2 relates to the ‘pericarp removal’ effect on transcriptome changes. However, 20M^−^ imbibed for 24 h at 9°C are amongst M^+^ samples on the PC2 axis (Area E, Figure 2). In particular, IND fruits under non-germination permissive conditions, form a relatively tight cluster in the negative PC1 and positive PC2 coordinates (Area F, Figure 2). Interestingly, during the imbibition time course for 20M^−^ and 25M^−^, the 24 h time point is farther along the PC1 axis positively than the later time points (100 h for 25M^−^, 75 h for 20M^−^), indicating the transcriptomes in the later time points resemble that of the dry-seed transcriptome more so than the earlier imbibition time points (Area B, Figure 2). Supporting this, more DEGs were found between the 24 h time points of 20M^−^ (4402) and dry 20M^−^ seed compared to the 75 h time point and dry 20M^−^ seed (3822), although there were fewer DEGs between the 24 h time points of 25M^−^ (3400) and dry 25M^−^ seed compared to the 100 h time point and dry 25M^−^ seed (3913) (Supplemental Data Set 1). Indeed, there is more variation than explained by only PC1 and PC2. While PC1 accounts for 25% and PC2 account for 14% of the variance (Figure 2), PC3 accounts for 11% of the variance and may have some relation to imbibition temperature (Supplemental Figure S3).

Despite 9°C imbibition permitting ca. 50% final germination, all imbibed 20IND fruit transcriptomes (24, 75, 125 h) clustered within Area F (Figure 2). Whereas 25IND fruits imbibed at 9°C for 24 h were also in Area F, 25IND fruit transcriptomes imbibed at 9°C for 75 h were in Area D together with the ‘germinating’ transcriptomes of M^+^ seed imbibed at 9°C and 14°C. Further, pericarp removal resulted in transcriptomes of M^−^ seed located in Area C at 75 h, indicating a strong effect of the pericarp on the 20IND fruit transcriptomes (Figure 2). Thus, it is evident that M^+^ and M^−^ differ in gene expression already in the dry state, and M^+^, M^−^, and IND differ more so following imbibition. Further, the transcriptomes are strongly influenced by imbibition temperature and pericarp removal. As a whole, the transcriptomes appear to reflect the status in terms of progress towards germination or dormancy, and the presence or absence of the pericarp in the case of seeds from IND fruits.

### Co-expressed gene modules in dry and imbibed seed transcriptomes associated with morph-specific germination responses

To further compare gene expression patterns between M^+^, M^−^, and IND at different imbibition temperatures associated with the regulation of germination progression, we grouped genes by their temperature-, time-, and morph-dependent expression patterns using weighted gene correlation network analysis (WGCNA) (Zhang and Horvath, 2005). This separated 11260 expressed genes into 11 modules each containing co-expressed genes (Figure 3A, Supplemental Figure S4, gene lists in Supplemental Data Set 2): black (523 genes); blue (1439); brown (1373); green (649); grey (2214); magenta (341); pink (365); purple (259); red (560); turquoise (2213); and yellow (1324). Figure 3A shows how neighboring genes in the PCA are clustered together and documents expression of genes in the modules during imbibition at 9°C.

**Figure 3.**
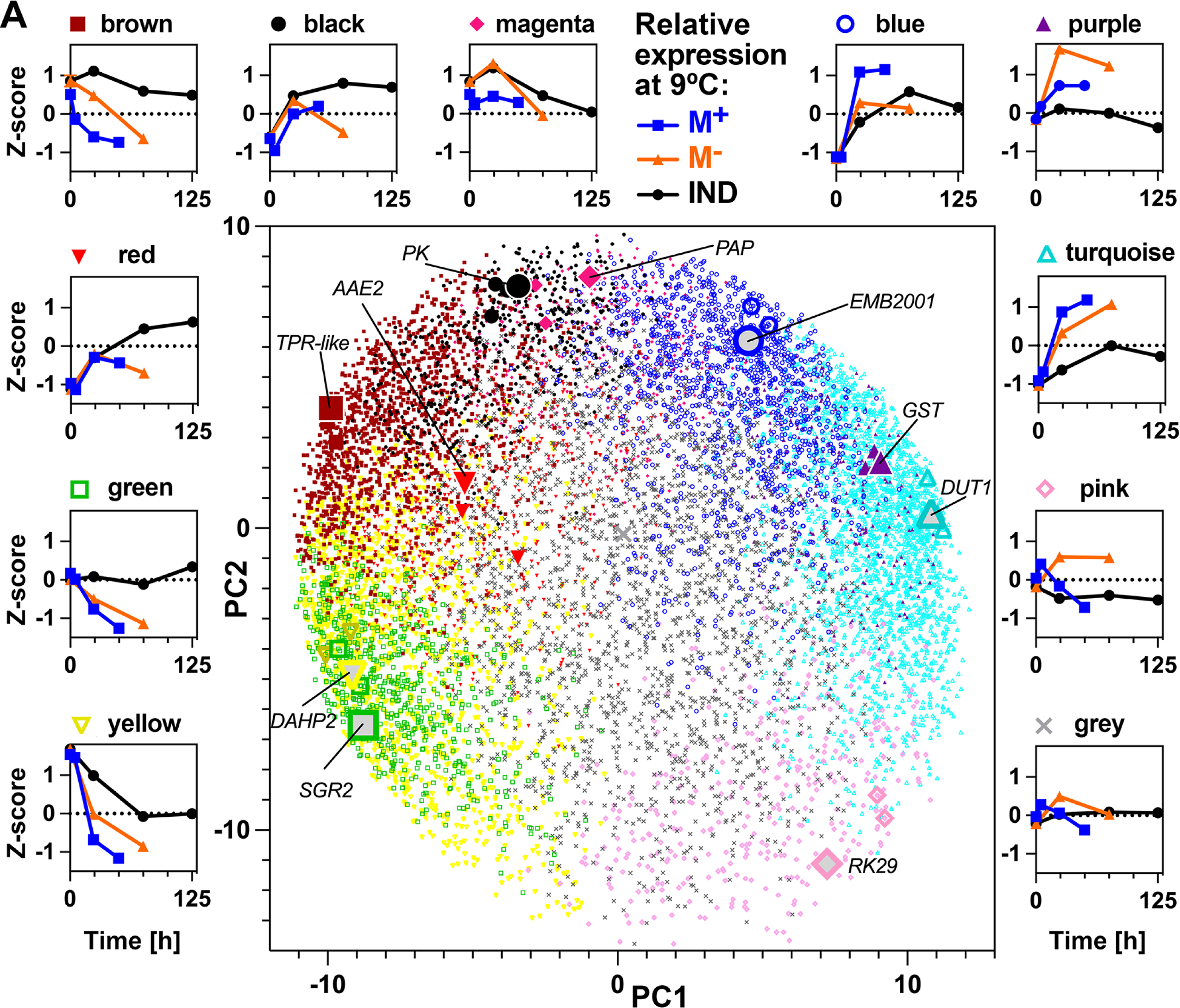
Weighted gene expression correlation network analysis (WGCNA) modules identified from dry and imbibed seed transcriptomes. WGCNA of 11260 genes identified eleven co-expressed gene modules, identified by color, across mature M^+^ and M^−^ seeds, and IND fruits harvested from plants at two different parental temperatures during reproduction (20 and 25°C) sampled in the dry state, and in the imbibed state at four different imbibition temperatures (9, 14, 20 and 24°C) at multiple time points. In the center, genes were separated by PCA of expression across all samples (first two principal components) and colored by module membership. Largest points indicate genes identified with the highest module membership for each module, labeled, and two additional large points representing high module membership candidates for the given module. Outer plots show mean Z-score expression of module genes during imbibition for M^+^ seeds, M^−^ seeds and IND fruits harvested from plants grown at 20°C and imbibed at 9°C. Expression of genes in modules for all samples is shown in Supplemental Figure S4.

Correlation between genes in the modules and associated PCs, temperatures, traits (morph, GR_50%_, G_max_), and quantified seed hormone contents facilitated investigation of potential roles of gene modules in morph-specific germination responses (Figure 4). Sample traits were clustered by their correlation patterns with module gene expression (using absolute values allowed positive and negative correlations to cluster together). Expression of purple and turquoise module genes was strongly positively correlated with GR_50%_ and G_max_. In contrast, expression of genes in the yellow, green, and red modules was strongly negatively correlated with GR_50%_ and G_max_. This suggested that expression of genes in the turquoise and purple modules supports a germination-promoting program. In contrast, expression of genes in the green, yellow, and red modules drives germination prevention or dormancy. Sample PC1 coordinates showed a similar trend reflecting the previously mentioned association between PC1 and germination. Seed ABA content, which showed the inverse pattern consistent with its negative association with germination, was highly positively correlated with yellow and green module gene expression and negatively correlated with blue, purple, and turquoise module gene expression. Brown and black module gene expression was highly positively correlated, and the pink module strongly negatively correlated with pericarp presence and PC2. Expression of yellow, green, and red module genes was also positively correlated with imbibition temperature and PC3 (reflecting the association previously mentioned), and purple and turquoise negatively correlated with imbibition temperature. This is consistent with the association between high temperatures and delayed germination. However, the overall correlation trends differed from those with G_max_ and GR_50%_ demonstrated by tree distance. This can be explained by differences, such as in magenta module gene expression, which was strongly negatively correlated with temperature, but not strongly correlated with germination traits. Parental temperature *per se* was not strongly correlated to any module eigengene.

**Figure 4.**
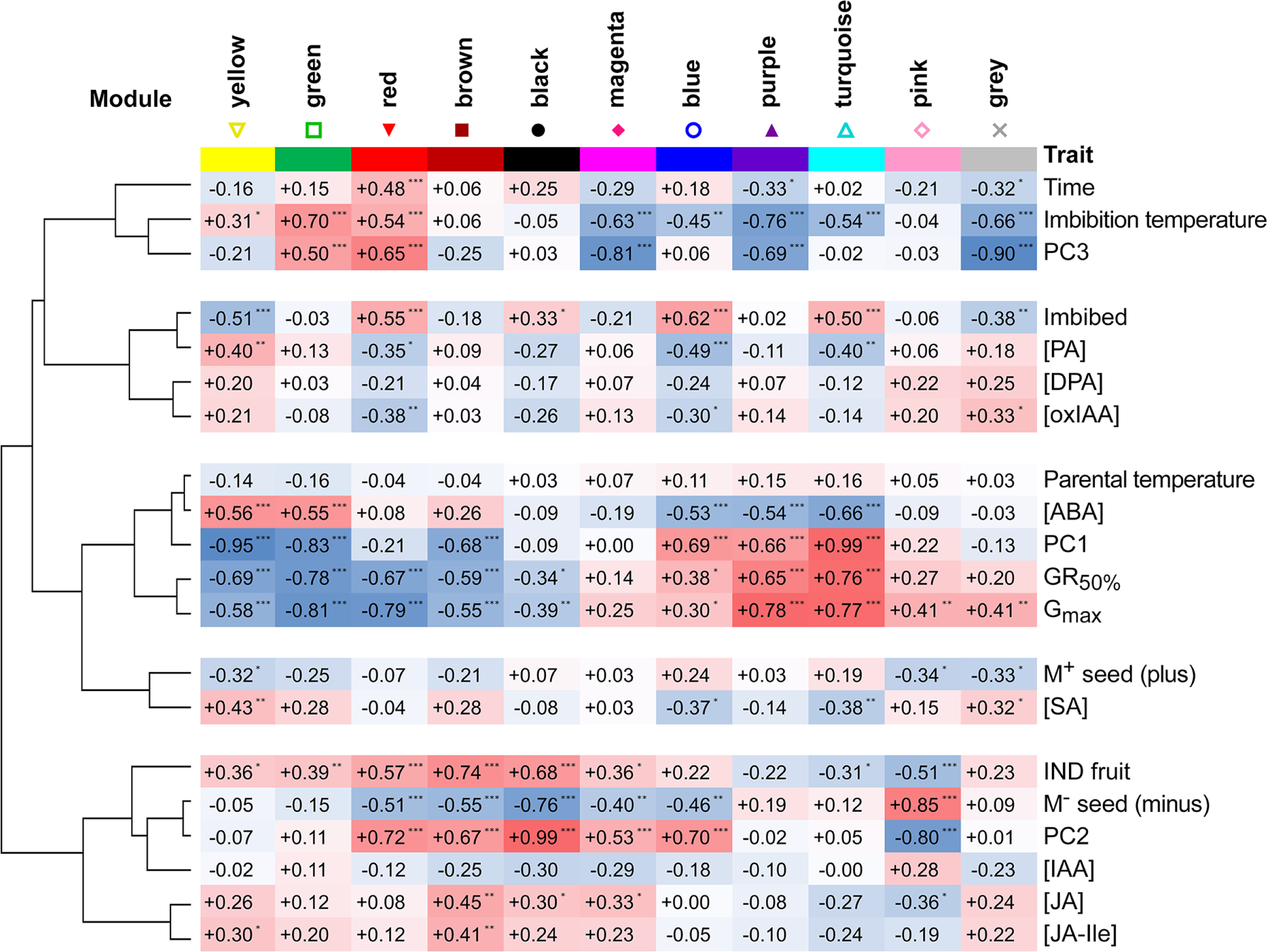
Correlation of WGCNA module expression with sample traits (hormone metabolites, PCA coordinates) and clustering. Hormone metabolites included are abscisic acid (ABA), ABA degradation products phaseic acid (PA) and dihydrophaseic acid (DPA), salicylic acid (SA), jasmonic acid (JA) and its isoleucine conjugate (JA-Ile), *cis*-(+)-12-oxophytodienoic acid (OPDA), indole-3-acetic acid (IAA) and its degradation product 2-ox-IAA (oxIAA). Sample PCA coordinates (PC1, PC2, PC3) were included as traits. Imbibition time, parental and imbibition temperature, GR_50%_ and G_max_ of samples were included. M^+^ and M^−^ seed, and indehiscent fruit (IND, pericarp presence) were included as binary variables (Plus, 0 or 1; Minus, 0 or 1; IND, 0 or 1). Asterisks indicate correlation significance: * – p < 0.05, ** – p < 0.01, *** – p < 0.0001) Correlation similarity tree was created using hierarchical clustering (1 – Pearson, average linkage using Morpheus, https://software.broadinstitute.org/morpheus).

We further investigated differences in module gene expression between specific sample pairs on a per module basis. Expression of genes in the black module was for example elevated in IND fruits and M^+^ seeds compared to M^−^ seeds, indicating it is associated with pericarp presence, but not germination kinetics *per se*, as expression in IND fruits and M^+^ seed is similar despite their contrasting germination kinetics. Expression within the brown module was elevated in IND fruits compared to both M^+^ and M^−^ at all imbibition temperatures, indicating a morph-specific and pericarp-dependent expression pattern associated with a delay in germination. Expression within the red module was strongly negatively correlated with germination, increased during imbibition under non-permissive germination conditions and tended to be more highly expressed in IND than M^−^ (for example during imbibition at 20°C). Expression within the green module was strongly correlated with conditions non-permissive for germination: higher in germination-delayed IND than in M^+^ and M^−^ germinating at 9°C and 14°C. Expression was high in all parental temperature × morphs at 20°C, perhaps except for 25M^+^, which did indeed germinate better relative to the other parental temperature × morphs at 20°C. Expression within the green module was higher in 20IND than 25IND at 9°C and 14°C correlating with strength of the pericarp-dependent delay of germination at these temperatures.

Genes within the yellow module were highly expressed in dry seeds compared to imbibed seeds and also strongly negatively correlated with germination (Figures 3A and 4). Gene expression decrease in the yellow module over time was delayed under conditions preventing germination and by the presence of the IND pericarp, more in 20IND than 25IND. Yellow module gene expression may be maintained or increased during prolonged inhibition of germination. For example, yellow module gene expression increased in M^−^ seeds, and perhaps in 20M^+^, imbibed at 20°C. Inverse to this pattern is the turquoise module where expression was strongly correlated with germination and repressed by the presence of the pericarp, especially in IND fruits from plants grown at 20°C. Expression within the pink module was strongly correlated with pericarp removal: highly expressed in M^−^ seed compared to IND fruit and M^+^ seed. Compared to other modules, genes in the grey module were more stably expressed across all treatments, but expression correlated positively with germination and negatively with imbibition and imbibition temperature. Genes in the magenta module were expressed more highly at lower than higher imbibition temperatures, and their expression was generally higher in IND than in M^+^ or M^−^ (apart from seeds/fruits from plants grown at 20°C and imbibed at 9°C). This module appears to be mostly associated with imbibition temperature.

Expression within the blue module was not generally strongly contrasting dependent on morph or pericarp removal, increased following imbibition, and was generally elevated during imbibition at lower temperatures. Expression within the purple module was strongly positively correlated with germination and negatively with imbibition temperature. Its expression was somewhat opposite of the green module, with high expression associated with germination permissive temperatures for M^+^, M^−^, and IND, and higher in 25IND than 20IND at 9°C and 14°C, also indicating negative association with pericarp-dependent delay of germination.

To gain further insight into which biological processes are associated with the promotion or delay of germination by morph, pericarp, imbibition temperature, or parental growth temperature, we performed Gene Ontology (GO) term enrichment analysis of the co-expressed gene modules (Supplemental Data Set 2). This revealed links between expression trends and module gene functions. For example, the GO terms ‘seed dormancy process’ was the most enriched in the green module, with ‘response to abscisic acid’ also the 24^th^ most enriched GO term. Other terms enriched in the green and yellow modules were also suggestive of dormancy, for example, ‘lipid storage’ and ‘chlorophyll catabolic process’. Conversely, ‘translation’ was the most enriched GO term in the turquoise module, with a number of cell-wall remodeling-related GO terms (e.g. ‘cell wall pectin metabolic process’, ‘plant-type cell wall organization’) and terms suggestive of increased metabolism, promotion of growth and transition to seedling highlighted in the turquoise and purple modules in which expression of the included genes was positively correlated with germination (e.g. ‘isopentenyl diphosphate biosynthetic process’, ‘methylerythritol 4-phosphate pathway’, ‘response to cytokinin’, ‘multidimensional cell growth’, ‘photosystem II assembly’, ‘gluconeogenesis’ and ‘glycolytic process’). Consistent with module gene functional enrichment and module gene expression correlation with ABA content (Figure 4), genes related to abscisic acid (ABA) biosynthesis are, for example, in the yellow, green, and brown module, for ABA degradation in the blue and grey module, and ABA receptor genes in the turquoise and purple module. ABA and cell-wall remodeling processes are discussed in more detail below.

In summary, out of these 11 gene modules (Figure 3), four are mainly associated with germination delay (brown, red, green, yellow); four are associated with germination stimulation (purple, turquoise, pink, grey); two are associated with imbibition temperature (grey, magenta); one associated mainly with imbibition (blue), four are associated with pericarp presence (black, brown, red, yellow), and one is associated with pericarp removal (pink).

### The role of the pericarp in altering abscisic acid metabolism and in morph-specific hormonal mechanisms to control dormancy and germination responses to temperature

While obviously many parameters contribute to the control of germination via modified gene expression patterns, the final “decision” depends to a large extend on the level and balance of several plant hormones in *A. thaliana* (Finch-Savage and Leubner-Metzger, 2006; Nambara et al., 2010; Linkies and Leubner-Metzger, 2012) and *Ae. arabicum* (Merai et al., 2019; Merai et al., 2023). We therefore quantified plant hormone metabolites using the same sampling scheme as for the RNAseq analysis (Supplemental Figure S1). In the IND fruit, the pericarp makes up 74.4% of the morph’s mass but at maturity does not contain living cells (Arshad et al., 2019; Arshad et al., 2020; Arshad et al., 2021). Transcript abundance patterns for mature dry and imbibed IND fruits therefore represent gene expression changes solely in the M^−^ seed (Lenser et al., 2016; Wilhelmsson et al., 2019; Arshad et al., 2021). The dead IND pericarp however contains hormone metabolites (Lenser et al., 2018) and we therefore quantified the hormone metabolites separately for the two fruit compartments (M^−^ seed and IND pericarp). The pericarp-imposed dormancy of 20IND fruits at 9°C and 14°C imbibition temperature was associated with abscisic acid (ABA) accumulation in 20M^−^ seeds during the imbibition of intact 20IND fruits (that is ABA content inside M^−^ seeds which were separated from the pericarp *after* imbibition at the times indicated) (Figure 5). In contrast, the ABA contents of 20M^+^ seeds and of bare 20M^−^ seeds (that is ABA content in imbibed M^−^ seeds which were separated from the pericarp *prior* to imbibition, i.e. in the dry state) steadily decreased upon imbibition at permissive temperatures.

**Figure 5.**
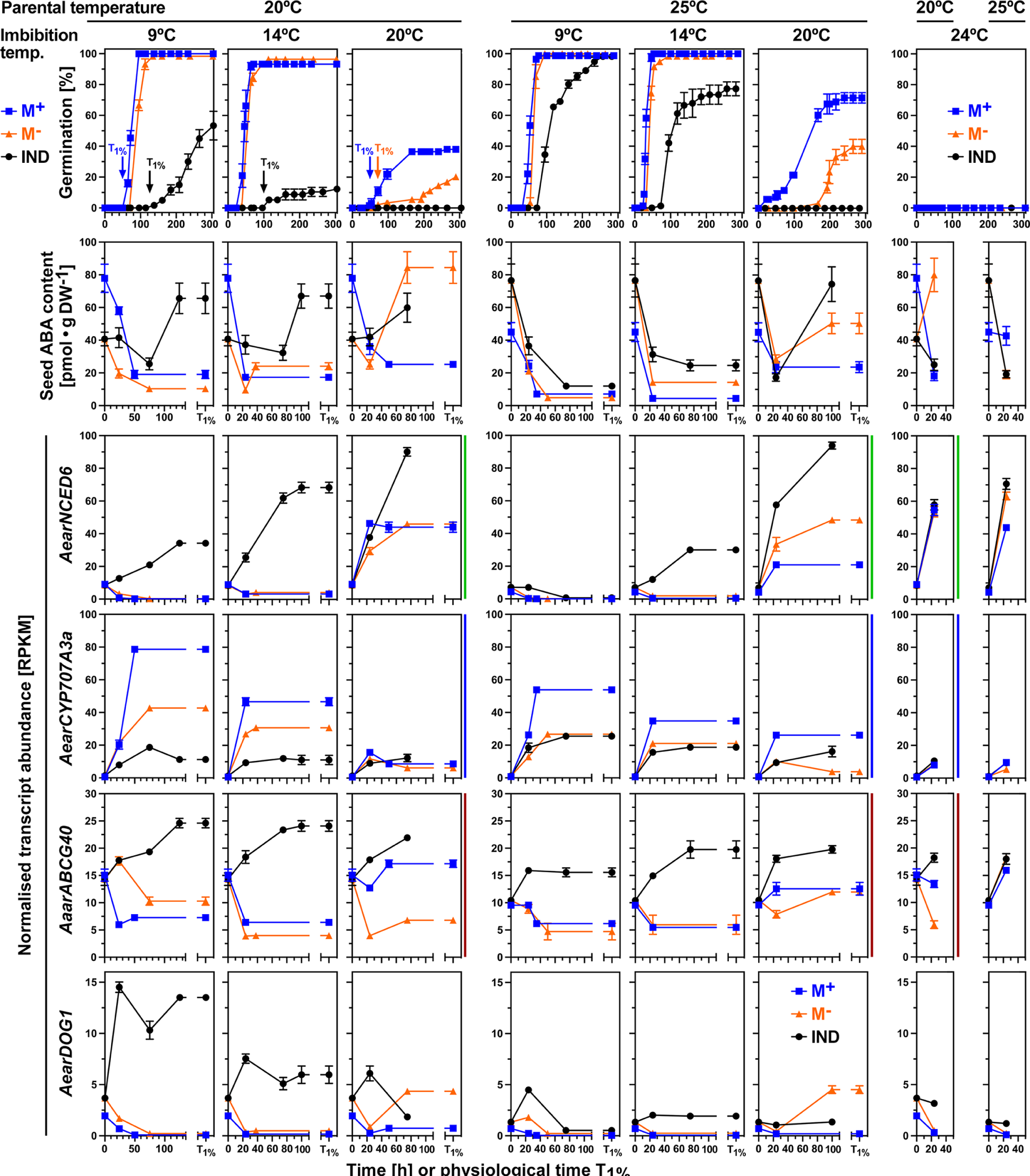
Comparative analysis of germination responses at different temperatures, associated abscisic acid (ABA) content and transcript abundance patterns of *Aethionema arabicum* dimorphic diaspores. Dimorphic diaspores (M^+^ seeds, IND fruits) and bare M^−^ seeds (extracted from IND fruits by pericarp removal) from two parental temperature regimes during reproduction (20°C versus 25°C) were compared for their germination kinetics, seed ABA contents (M^+^ seeds, bare M^−^ seeds, and M^−^ seeds encased inside the imbibed IND fruit) and seed transcript abundance patterns at four different imbibition temperatures (9, 14, 20 and 24°C). Comparative results were obtained for physical (in hours) and physiological time points (T_1%_, representing the population’s onset of germination completion). Normalized transcript abundances in reads per kilobase per million (RPKM) from the transcriptomes (RNA-seq) are presented for the ABA biosynthesis 9-*cis*-epoxycarotenoid dioxygenase gene *AearNCED6*, the ABA 8′-hydroxylase gene *AearCYP707A3*, the plasma membrane ABA uptake transporter gene *AearABCG40*, and the *Delay of germination 1* dormancy gene *AearDOG1*. WGCNA modules (Figure 3) for these genes are indicated by the vertical color lines next to the graphs. For *Ae. arabicum* gene names and IDs see Supplemental Table S2 or the Expression Atlas (https://plantcode.cup.uni-freiburg.de/aetar_db/index.php); for RNAseq single values see the Expression Atlas or Supplemental Data Set 1. Mean ± SEM values of 3 (germination, RNA-seq) or 5 (ABA) replicates each with 20 (germination), 30-40 (ABA) and 60-80 (RNA-seq) seeds are presented.

In agreement with the high ABA content in seeds, transcript abundance for *AearNCED6* (9-*cis*-epoxycarotenoid dioxygenase), a key gene in ABA biosynthesis, increased in the seeds of imbibed 20IND fruits, and decreased in 20M^+^ and bare 20M^−^ seeds upon imbibition at 9°C and 14°C (Figure 5). A similar expression pattern was evident for other ABA biosynthesis genes (Supplemental Figure S5). Consistent with a role of parental temperature, the pericarp-imposed dormancy was reduced in 25IND as compared to 20IND fruits, and the ABA contents declined in the seeds of imbibed 25IND fruits, as well as in 25M^+^ and bare 25M^−^ seeds upon imbibition at 9°C and 14°C (Figure 5). Despite this decline, the ABA content in seeds of imbibed 25IND fruits remained higher compared to 25M^+^ and 25M^−^ seeds. The observed increase of transcript abundance for *AearNCED6* and other ABA biosynthesis genes at 9°C and 14°C imbibition temperature was somewhat reduced in 25IND compared to 20IND fruits (Figure 5, Supplemental Figure S5). At the non-permissive imbibition temperatures 20°C and 24°C for 20IND and 25IND germination, transcripts for *AearNCED6* and other ABA biosynthesis genes accumulated most strongly in seeds of imbibed IND fruits, also somewhat in bare M^−^ seeds, but not in M^+^ seeds. At 20°C imbibition temperature, the ABA content also increased in seeds of imbibed IND fruits and in bare M^−^ seeds, but not in M^+^ seeds. Taken together, these findings suggest that ABA accumulation due to *de novo* ABA biosynthesis by *AearNCED6* and other ABA biosynthesis genes explains, at least in part, the distinct responses of the morphs to parental and imbibition temperatures, revealing that germination inhibition by elevated ABA levels is a key mechanism of the pericarp-imposed dormancy in 20IND fruits.

In further support of the importance of ABA in the control of the pericarp-imposed dormancy, the enhanced degradation of ABA in the M^+^ seed morph upon imbibition was associated with increased transcript abundances for *AearCYP707A3* and other ABA 8′-hydroxylases, while their expression remained low in the corresponding IND fruit morph (Figure 5, Supplemental Figure S5). Therefore, the expression patterns of *AearCYP707A3* (highest in M^+^ seeds, lowest in IND fruits, intermediate in M^−^ seeds) were, in most cases, inverse to the *AearNCED6* expression patterns. Further to this, the expression patterns for gibberellin biosynthesis (GA3-oxidases) and inactivation (GA2-oxidases) genes were inverse to the ABA biosynthesis (NCED) and inactivation (CYP707A) genes (Supplemental Figure S5D). In addition to ABA metabolism, the presence of the pericarp also enhanced the transcript accumulation for the plasma membrane ABA uptake transporter gene *AearABCG40* (an ABC transporter of the G subfamily) and the *AearDOG1* (*Delay of germination 1*) dormancy gene in a morph-specific and temperature-dependent manner (Figure 5).

Hormone metabolite contents per pericarp of ABA, its 8′-hydroxylase pathway breakdown compounds phaseic acid (PA) and dihydrophaseic acid (DPA), as well as for jasmonic acid (JA), its bioactive isoleucine conjugate (JA-Ile) and for salicylic acid (SA), were in general 10 to 20 fold higher in the dry state declined rapidely in the pericarp upon imbibition at any temperature (Figure 6A; Supplemental Figure S6). In contrast to these hormone metabolites, the contents per pericarp of *cis*-(+)-12-oxophytodienoic acid (OPDA), which is an oxylipin signaling molecule and JA biosynthesis precursor (Linkies and Leubner-Metzger, 2012; Dave et al., 2016), did not decline during imbibition and its contents in the pericarp remained much higher compared to the encased M^−^ seed (Supplemental Figure S6B,C). When the ABA, PA, DPA, JA, JA-Ile, SA, and OPDA contents of diaspore compartment (seed *versus* pericarp) were compared, they were, in general, >20 times (dry state) and >5 times (imbibed state) higher higher in the pericarp compared to the M^−^ seed extracted from the IND fruit (Figure 6A; Supplemental Figure S6). An exception from this was ABA in 20IND fruits where the contents per compartment (pericarp *versus* encased M^−^ seed) during late imbibition at 9°C and 14°C were roughly equal, but in 25IND fruits ABA was higher in the pericarp compared to the encased M^−^ seed also during imbibition (Figure 6A; Supplemental Figure S6C). Although SA declined rapidely in the pericarp during imbibition, its contents remained much higher in the pericarp also during late imbibition as compared to the encased M^−^ seed. Further to this comparison (pericarp *versus* encased M^−^ seed), the hormone metabolite contents of imbibed bare M^−^ seeds and imbibed M^+^ seeds were always lower compared to IND fruits (Figures 6A; Supplemental Figure S6). The contents of the auxin indole-3-acetic acid (IAA) were low in M^+^ and M^−^ seeds. IAA was below limit of detection in pericarp tissue, but significant amounts of the major IAA degradation product 2-oxoindole-3-acetic acid (oxIAA) were detected (Supplemental Figure S6), suggesting that IAA degradation occurred during the late stages of pericarp development.

**Figure 6.**
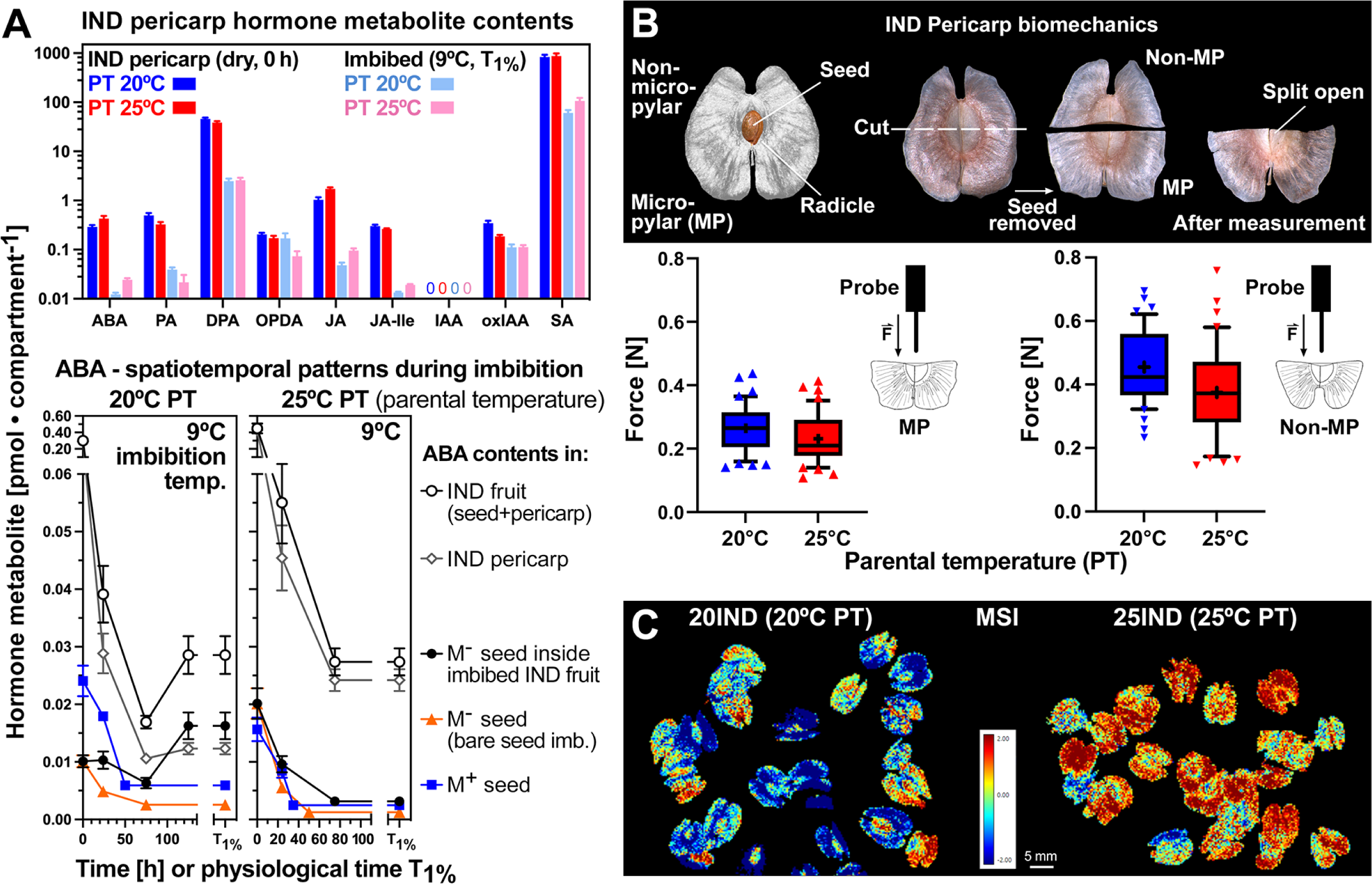
The effect of parental temperature (PT) on the biochemical and biomechanical properties of the IND pericarp and the pericarp-imposed dormancy of *Aethionema arabicum*. A, Comparative analysis of hormone metabolite contents in IND pericarps, M^+^ and M^−^ seeds from two parental temperature regimes (20°C versus 25°C) in the dry state and for ABA during imbibition at 9°C (see Supplemental Figure S6 for other imbibition temperatures and other hormone metabolites). Hormone metabolites presented: abscisic acid (ABA) and ABA degradation products phaseic acid (PA) and dihydrophaseic acid (DPA), salicylic acid (SA), jasmonic acid (JA) and its isoleucine conjugate (JA-Ile), *cis*-(+)-12-oxophytodienoic acid (OPDA). Mean ± SEM values of 5 (hormone metabolites) biological replicate samples are presented. B, The effect of parental temperature during reproduction on the IND pericarp resistance quantified by biomechanical analysis. Results are presented as box plots, whiskers are drawn down to the 10^th^ percentile and up to the 90^th^ (mean is indicated by ‘+’), n = 42. The micropylar (where the radicle emerges during fruit germination) pericarp half grown at 20°C shows a slightly higher tissue resistance versus 25°C (p = 0.047). The non-micropylar half has a higher tissue resistance whilst not showing any difference between 20IND and 25IND; see Supplemental Figure S8 for extended biomechanical properties. C, The effect of parental temperature on the IND pericarp biochemical composition as analyzed by multispectral imaging (MSI).

Leachates of inhibitors from pericarp tissue, including ABA, OPDA, and phenolic compounds, may inhibit germination and thereby contribute to coat dormancy (Ignatz et al., 2019; Mohammed et al., 2019; Grafi, 2020). In agreement with this, pericarp extracts (PE) and ABA both delayed the germination of bare M^−^ seeds (Supplemental Figure S7A). PE application explained the delayed 20IND fruit germination only partially, but the delay could be fully mimicked by exposure to 5 µM ABA (Supplemental Figure S6A). Treatment of M^−^ seeds with PE delayed their germination, concordant to the delay of 25IND fruit germination at 9°C imbibition (Supplemental Figure S7C). In contrast to PE and ABA, treatment of seeds with SA, OPDA, JA, or JA-Ile did not appreciably affect seed germination (Supplemental Figure S6D), and PE from 20IND and 25IND pericarps did not differ in their inhibitory effects (Supplemental Figure S7C-D). To investigate if PE and ABA treatment of bare M^−^ seeds can mimic the pericarp effect on gene expression as observed in imbibed IND fruits, we conducted RT-qPCR of representative genes for each WGCNA module. In several cases, the ABA treatment indeed mimicked the PE effect (Supplemental Figure S7B), but neither PE nor ABA could fully mimic the effect of the intact pericarp. We, therefore, conclude that leaching of ABA or other inhibitors from the pericarp is not the major component by which the pericarp exerts its effects on gene expression and germination responses.

Pericarp properties are known to affect embryo ABA sensitivity, oxygen availability, and the biomechanics of germination (Benech-Arnold et al., 2006; Hoang et al., 2013; Steinbrecher and Leubner-Metzger, 2017). Biomechanical analysis revealed that the tissue resistance at the micropylar pericarp (where the radicle emerges during germination of fruit-enclosed seeds) was slightly higher in 20IND as compared to 25IND pericarps (Figure 6B and Supplemental Figure S8). Tissue resistance at the non-micropylar pericarp was higher and did not differ between 20IND and 25IND. Parental temperature is also known to affect dormancy via the seed coat pro-anthocyanidin content. Multispectral imaging (MSI) can visualize this and unknown differences in the biochemical composition of seed coats that affect reflectance spectra (Penfield and MacGregor, 2017). Figure 6C shows that MSI detected unknown differences in the biochemical composition between 20IND and 25IND fruits. The distinct parental temperatures therefore affected pericarp development leading to distinct biochemical compositions upon maturity. These differences in 20IND and 25IND pericarp biomechanics and biochemistry may also be associated with altered pericarp oxygen permeability.

### Evidence for pericarp-mediated hypoxia, morph-specific transcription factor expression, and ABA signaling in the control of *Ae. arabicum* dormancy and germination

To identify transcription factor (TF) genes and corresponding target *cis*-regulatory elements (TF binding motifs/sites), we conducted enrichment analyses for each WGCNA module described above. Enriched motifs from the ArabidopsisDAPv1 database (O’Malley et al., 2016) for each module are compiled in Supplemental Data Set 3. The chord diagram (Figure 7A) shows identified *Ae. arabicum* TF genes in each WGCNA module and their connection to corresponding target *cis*-regulatory motifs. For example, for ABA-related bZIP TFs (Nambara et al., 2010) such as ABA Insensitive 5 (ABI5, green module), ABA-responsive element (ABRE)-binding proteins or ABRE-binding factors (ABFs) such as AREB3 (yellow module), G-box binding factors (GBFs) such as GBF3 (red module), motifs were enriched in the green module (Figure 7A, Supplemental Data Set 3). Another example are hypoxia-related TFs of the ethylene response factor (ERF) group VII that are known to induce gene expression upon hypoxia by binding to motifs such as the ERF73 *cis*-regulatory element and the hypoxia-responsive promoter element (HRPE) of hypoxia-responsive genes (Gasch et al., 2016). In *A. thaliana* the ERF group VII has five members: ERF73/HRE1 (HYPOXIA RESPONSIVE ERF1), ERF71/HRE2, RAP2.2 (RELATED TO APETALA2.2), RAP2.3, and RAP2.12. A comparison of several Brassicaceae genomes, including *A. thaliana* and *Ae. arabicum*, irevealed a high number of conserved noncoding sequences (Haudry et al, 2013). The ABRE and HRPE motifs and the TFs binding to these *cis*-regulatory elements, are enriched in promoters of hypoxia-responsive and ABA-responsive genes and widely conserved among multiple species (Gasch et al. 2016; Gomez-Porras et al., 2007; O’Malley et al. 2016). Their roles in inducing their hypoxia-responsive target genes have been well investigated for the fermentation enzymes alcohol dehydrogenase (ADH) and pyruvate dehydrogenase (PDC) (Kürsteiner et al., 2003; Yang et al., 2011; Papdi et al., 2015; Gasch et al., 2016; Seok et al., 2022). In *Ae. arabicum* ERF73 and HRPE motifs were enriched in the grey, yellow, and brown modules, and putative target genes *AearADH1a* and *AearPDC2* are gene members of these modules (Figure 7A, Supplemental Data Set 3). In contrast to *A. thaliana* which has *AtERF71* and *AtERF73* as two highly related group VII ERF genes, there is only one homolog in *Ae. arabicum*, member of the grey module, which we named *AearERF71/73* (*AearHRE1/2*) as its sequence is equally homologous to either of the two *A. thaliana* genes. Also, in contrast to *A. thaliana*, which has only one ADH gene (*AtADH1*), there are two ADH genes in *Ae. arabicum*, namely *AearADH1a* (brown module) and *AearADH1b* (green module) (Figure 7A).

**Figure 7.**
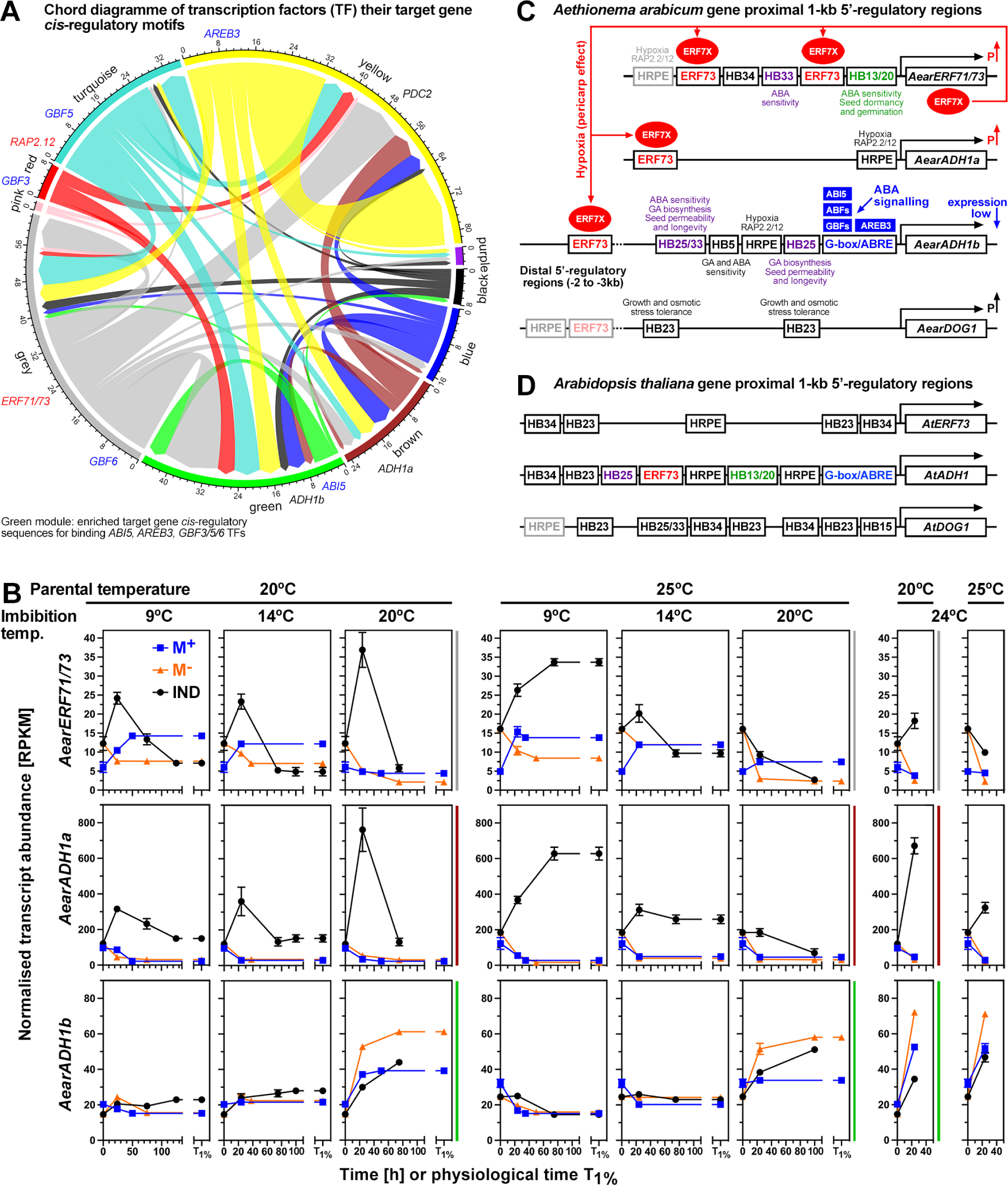
Transcription factor (TF) and target *cis*-regulatory motif analysis of *Aethionema arabicum* gene expression with focus on hypoxia and ABA related genes. A, Chord diagram of identified TFs and their target genes in the WGCNA modules. Examples for TFs (*red* or *blue*) and their target genes (*black*). B, Transcript abundance patterns (RNA-seq) of the *Ae. arabicum Hypoxia responsive ERF* (*AearERF71/73*) TF and the alcohol dehydrogenase genes *AearADH1a* and *AearADH1b* in seeds of imbibed dimorphic diaspores (M^+^ seeds, IND fruits) and bare M^−^ seeds from two parental temperature regimes (20°C versus 25°C) at four different imbibition temperatures (9, 14, 20 and 24°C). WGCNA modules (Figure 3) for these genes are indicated by the vertical color lines next to the graphs. Mean ± SEM values of 3 replicates each with 60-80 seeds are presented. C, Hypothethical working model for the pericarp-mediated hypoxia up-regulation (P↑) and ABA signaling, and *cis*-regulatory motifs for the *Ae. arabicum ERF71/73*, *ADH1a, ADH1b*, and *DOG1* genes. Promoter motifs indicated include the hypoxia-responsive promoter element HRPE, the G-box and ABA-responsive element (ABRE), the ERF73 *cis*-regulatory element and HB-motifs for the binding of homeobox TFs (for details see Supplemental Figure S10). These motifs are the targets for the AearERF71/73 TF (ERF7X, *red ellipse*) and the ABA related ABI5, ABF (ABRE-binding factors), GBF (G-box-binding factors), and AREB3 TFs (*blue boxes*). D, Comparative analysis of the corresponding *Arabidopsis thaliana AtERF73*, *AtADH1* and *AtDOG1* gene 5’-regulatory regions (for details see Supplemental Figure S10). Note that *A. thaliana* has only one while *Ae. arabicum* has two ADH genes; see Supplemental Figure S9 for other fermentation-related genes. For *Ae. arabicum* gene names and IDs see Supplemental Table S2 or the Expression Atlas (https://plantcode.cup.uni-freiburg.de/aetar_db/index.php); for RNAseq single values see the Expression Atlas or Supplemental Data Set 1.

Figure 7B shows that *AearERF71/73* and *AearADH1a* transcripts accumulated in *Ae. arabicum* IND fruits upon imbibition, while their transcript abundances in M^−^ and M^+^ seeds remained low. *AearPDC2* had a similar expression pattern in that the transcript abundances were highest in IND fruits (Supplemental Figure S9). In contrast to *AearADH1a* and *AearPDC2*, the expression of *AearADH1b* (the second *Ae. arabicum* ADH gene) and *AearPDC1* remained comparatively low, and *AearLDH* (lactate dehydrogenase) was less consistently elevated in imbibed IND fruits compared to M^−^ and M^+^ seeds (Figure 7B; Supplemental Figure S9). Taken together, this suggested that hypoxia conferred to IND fruits by the pericarp may lead to the induction of the ethanolic fermentation pathway with *AearADH1a* and *AearPDC2* as hypoxia-responsive target genes (Figure 7C), as it is known for the hypoxia-response of the *AtADH1* and *AtPDC1* genes (Kürsteiner et al., 2003; Yang et al., 2011; Papdi et al., 2015; Gasch et al., 2016; Seok et al., 2022). To investigate the ABA and hypoxia-regulated expression further, we compared the *Ae. arabicum* and *A. thaliana ADH*, *PDC*, *ERF71/73*, *LDH,* and *DOG1* genes for *cis*-regulatory binding motifs using FIMO (Figure 7C, D; Supplemental Figures S9, S10; Supplemental Data Set 4). The focus of this analysis was on the widely conserved general HRPE, ABRE, ERF73, and binding motifs for Homeobox (HB) TFs (see Supplemental Figure S10A for best possible matches of *cis*-regulatory binding motifs in *Ae. arabicum* genes). The binding motifs for HB TF were included in this analysis because they are known to control seed-to-seedling phase transition, seed ABA sensitivity, dormancy, longevity and embryo growth in *A. thaliana* (Barrero et al., 2010; Wang et al., 2011; Bueso et al., 2014; Silva et al., 2016; Stamm et al., 2017). The *AearADH1a* and *AearPDC2* 5’-regulatory gene region contain ERF73 and HRPE motifs and are distinct from the *AtADH1, AearPDC1* and *AearADH1b* 5’-regulatory gene regions in that they don’t contain G-box/ABRE motifs (Figure 7C, Supplemental Figure S9). The *AearERF71/73* 5’-regulatory gene region was also distinct from its *A. thaliana* homologs by the presence of two ERF73 motifs, suggesting that the *AearERF71/73* gene possibly provides a positive feedback regulation on the pericarp/hypoxia-mediated *AearADH1a, AearPDC2* and *AearERF71/73* expression (Figure 7C). Further details of this hypothetical working model are described and discussed in Supplemental Figure S9. In contrast to the ethanolic fermentation pathway (PDC-ADH, substrate pyruvate), which is up-regulated in IND fruits (Figure 7, Supplemental Figure S9), the seed-specific ‘Perl’s pathway,’ which controls pyruvate production (Weitbrecht et al., 2011), is down-regulated in IND fruits as compared to bare M^−^ seeds (Supplemental Figure S11).

To test if *AearERF71/73*, *AearADH1a* and *AearPDC2* and other candidate genes are indeed regulated by hypoxia we analyzed their expression in bare M^−^ seeds imbibed under hypoxia conditions (Figure 8). Gasch et al. (2016) identified a list of 49 core hypoxia-responsive genes in *A. thaliana* seedlings which includes *AtERF73*, *AtADH1* and *AtPDC2*. Of these 49 core hypoxia-responsive *A. thaliana* genes, we identified from the transcriptome analysis that expression of 14 of 41 putative *Ae. arabicum* orthologs was elevated in IND fruits, whereas their expression in M^−^ and M^+^ seeds remained low. Examples are presented in Supplemental Figure S12 and include *AearHRA1*, *AearETR2, AearNAC102*, *AearJAZ3*, *AearHHO2*, and other *Ae. arabicum* homologs from the core list of *A. thaliana* hypoxia-responsive genes (Christianson et al., 2009; Gasch et al., 2016; Ju et al., 2019). Figure 8A shows that the germination of bare M^−^ seeds is indeed severely delayed under hypoxia (4.5% oxygen) as compared to normoxia (21% oxygen) conditions. This resulted in the hypoxia-mediated induction of the *AearERF71/73*, *AearADH1a*, *AearPDC2*, *AearHRA1*, *AearETR2, AearJAZ3*, *AearNAC102*, *AearHHO2*, and other genes (Figure 8A, Supplemental Figure S13A). Hypoxia delayed the germination of bare M^−^ seeds similar to the pericarp in IND fruits, in both cases the T_1%_ was ca. 100 h (Supplemental Figure S13B). M^−^ seed germination was also delayed by 5 µM ABA, and the combined treatment (hypoxia+ABA) had a stronger inhibitory effect on germination (Figure 8A).

**Figure 8.**
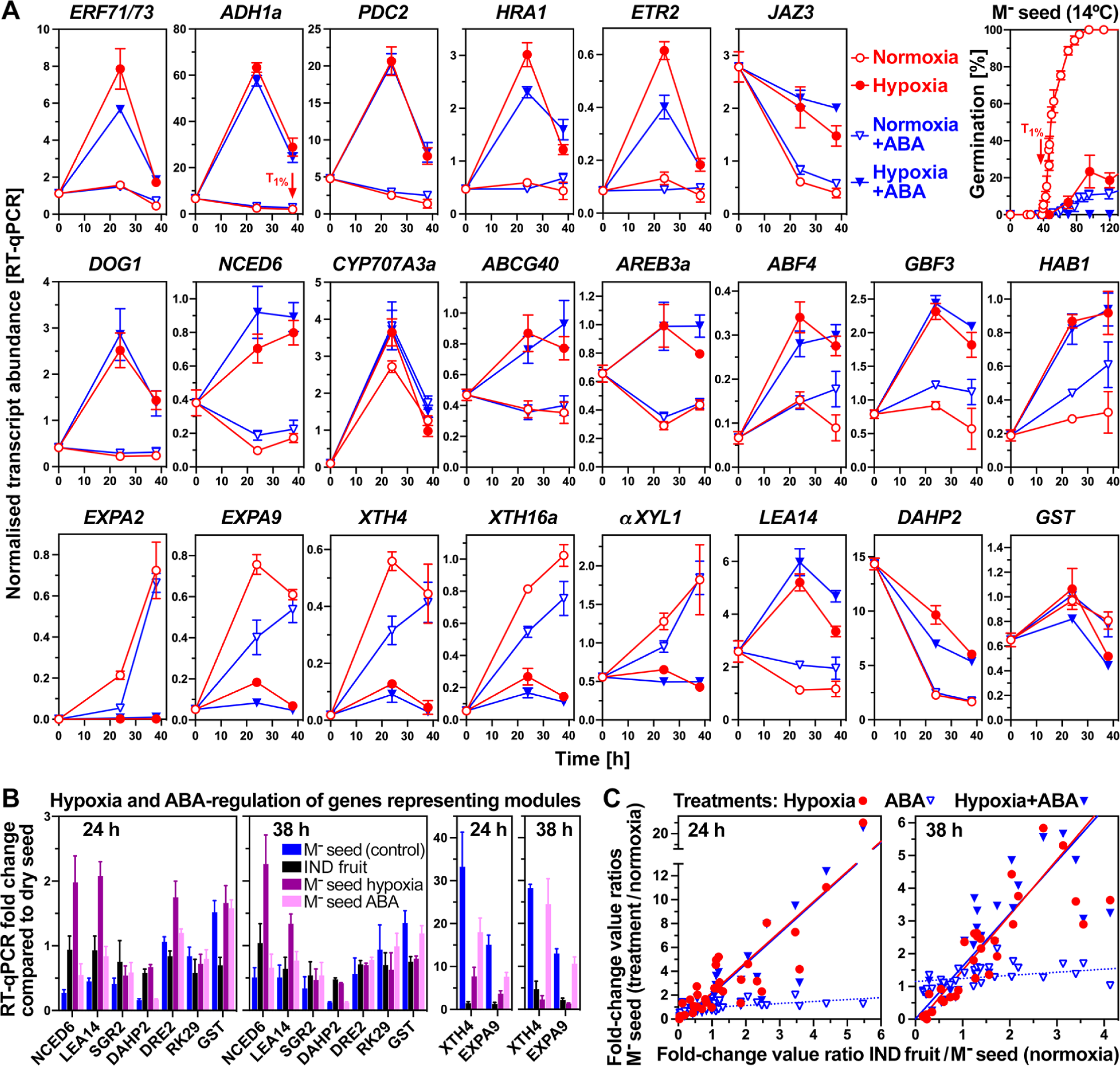
The effect of hypoxia and ABA on germination and gene expression of bare M^−^ seeds. A, RT-pPCR expression analysis of selected genes during *Aethionema arabicum* bare M^−^ seed imbibition under hypoxia (4.5±0.2% oxygen) and normoxia (21% oxygen) conditions ± 5 µM abscisic acid (ABA). Bare M^−^ seeds were obtained from dry IND fruits by pericarp removal and imbibed at 14°C in continuous light. The 38 h timepoint (arrow) corresponds to T_1%_ of the control (normoxia without ABA). For additional genes and expression in IND fruits see Supplemental Figure S12. B, RT-pPCR expression analysis of genes representing the WGCNA modules used in Supplemental Figure S7 to investigate the effects of pericarp extract. C, Correlation analysis between the effects of the pericarp (IND fruits), hypoxia (M^−^ seeds) and ABA (M^−^ seeds) on the expression of 32 genes as compared to M^−^ seeds in normoxia (control). ‘Treatments / control’ ratios (y-axis) of fold-change values (from the dry state to 24 h or 38 h) were calculated and plotted agains the ‘IND fruit / control’ ratios (x-axis). Linear regression lines indicate strong linear relationships for hypoxia versus pericarp (R^2^ 0.79 and 0.70 for 38 h and 24 h, respectively) and for hypoxia+ABA versus pericarp (R^2^ 0.80 and 0.75), but not for ABA versus pericarp (R^2^ 0.16 and 0.30). Mean ± SEM values of 3 (germination, RT-qPCR) biological replicate samples are presented. For *Ae. arabicum* gene names and IDs see Supplemental Table S2 or the Expression Atlas (https://plantcode.cup.uni-freiburg.de/aetar_db/index.php).

The importance of ABA signaling in the control of *Ae. arabicum* pericarp-imposed dormancy of IND fruits was evident in the expression patterns of ABRE-binding (ABI5, AREBs/ABFs) and G-box-binding (GBFs) bZIP TF genes. Transcripts of *AearAREB3a*, *AearAREB3b*, *AearABI5*, *AearABF1*, *AearABF2*, *AearABF3*, *AearABF4*, *AearGBF1*, *AearGBF2*, *AearGBF3,* and *AearGBF4* were up-regulated in M^−^ seeds inside IND fruits, and in general expressed lowly in isolated M^−^ seeds and in M^+^ seeds (Figure 9; Supplemental Figure S14A). By contrast, *AearGBF5*, *AearRAP2.12*, and the transcript abundances of several HB TF including *AearHB13* were down-regulated in M^−^ seeds inside IND fruits (Figure 9, Supplemental Figure S14B). In *A. thaliana,* these bZIP TFs are also known to control the ABA-related expression, including for the *AtADH1* gene, by binding to G-box and ABRE 5’-regulatory motifs (Lu et al., 1996; Gomez-Porras et al., 2007; Nambara et al., 2010; Yoshida et al., 2010; O’Malley et al., 2016). HB13 and HB20 TFs constitute node-regulators within the co-expression network controlling seed-to-seedling phase transition (Silva et al., 2016) while other HB TFs control seed ABA sensitivity, dormancy, longevity and embryo growth (Barrero et al., 2010; Wang et al., 2011; Bueso et al., 2014; Stamm et al., 2017; Renard et al., 2021). Transcript abundance patterns of ABA signaling components including for the protein phosphatase 2C protein HAB1 (Nambara et al., 2010) also exhibit pericarp-affected expression patterns in the *Ae. arabicum* morphs (Supplemental Figure S14C). Figure 8A shows that in bare M^−^ seeds imbibed under hypoxia, many ABA-related genes and the dormancy gene *AearDOG1* are up-regulated by hypoxia. In contrast to this, hypoxia or the presence of the pericarp caused down-regulation of genes encoding cell wall-remodeling proteins (see next section). Expression of components of the general RNA polymerase II transcription elongation complex, ribosomal proteins, and 20S proteasome subunits differed during *Ae. arabicum* fruit morph development (Wilhelmsson et al., 2019). Related *Ae. arabicum* genes, especially of the turquoise, purple, and pink WGCNA modules, exhibited distinct pericarp-affected expression patterns (Supplemental Figure S15), which persisted throughout imbibition.

**Figure 9.**
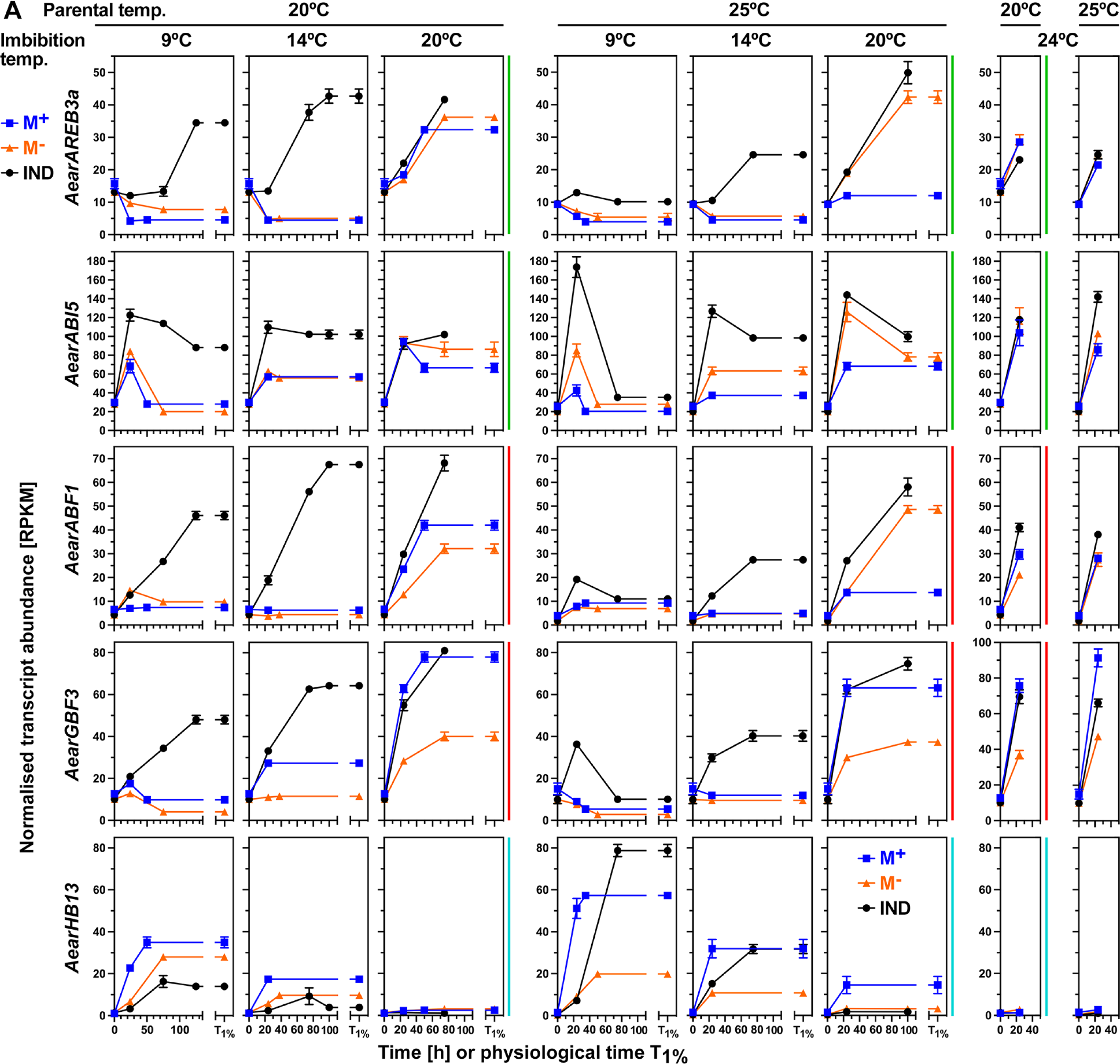
Transcript abundance patterns (RNA-seq) of *Aethionema arabicum* ABA-related and homeobox (HB) TF genes. Results for *AearAREB3a*, *AearABI5*, *AearABF1*, *AearGBF3*, and *AearHB13* transcript abundances in seeds of imbibed dimorphic diaspores (M^+^ seeds, IND fruits) and bare M^−^ seeds (extracted from IND fruits) from two parental temperature regimes (20°C versus 25°C) at four different imbibition temperatures (9, 14, 20 and 24°C) are presented (see Supplemental Figure S14 for other ABF, GBF and HB TFs). WGCNA modules (Figure 3) are indicated by the vertical color lines next to the graphs. For *Ae. arabicum* gene names and IDs see Supplemental Table S2 or the Expression Atlas (https://plantcode.cup.uni-freiburg.de/aetar_db/index.php); for RNAseq single values see the Expression Atlas or Supplemental Data Set 1. Mean ± SEM values of 3 replicates each with 60-80 seeds are presented.

### The role of morph-specific and temperature-dependent expression patterns of cell wall-remodeling genes for *Ae. arabicum* germination and dormancy responses

Cell wall-remodeling by expansins, and enzymes targeting xyloglucans, pectins, and other cell wall components are required for successful embryo growth and for restraint weakening of covering structures in seed and fruit biology (Graeber et al., 2014; Shigeyama et al., 2016; Steinbrecher and Leubner-Metzger, 2017; Arshad et al., 2021; Steinbrecher and Leubner-Metzger, 2022). The expression ratio of expansin genes between M^−^ seed within IND fruit and bare M^−^ seed at 24 h or T_1%_ remained very low at any imbibition temperature (Figure 10A; Supplemental Figure S16A). Consistent with this, transcripts of all *Ae. arabicum* type expansins (Figure 10C; Supplemental Figure S16C) were only induced in M^+^ and bare M^−^ seeds, but not appreciably in imbibed IND fruits. Similar results were obtained for xyloglucan endotransglycosylases/hydrolases (XTHs) for 20IND fruits, whereas considerable induction was observed for 25IND fruits at the permissive imbibition temperatures (9°C and 14°C). In addition to XTHs, xyloglucan remodeling is achieved by a battery of bond-specific transferases and hydrolases (Figure 10B), and in the *Ae. arabicum* transcriptomes, most of them belong to the turquoise WGCNA module with *αXYL1* as an example (Figure 10). Higher transcript expression in bare 20M^−^ seeds as compared to 20IND fruits was observed for *αFUCs, βGALs*, *βXYL,* and *GATs* (Supplemental Figure S16D), suggesting that the pericarp-mediated repression and the resultant reduction in xyloglucan remodeling is part of the 20IND dormancy mechanism. The induction in 25IND fruits that eventually germinate further supports the importance of these genes and their products in the germination process. Expression comparison of M^+^ seeds and isolated M^−^ seeds further confirm that the presence of the pericarp is the most important factor for the expression differences between the dimorphic diaspores.

**Figure 10.**
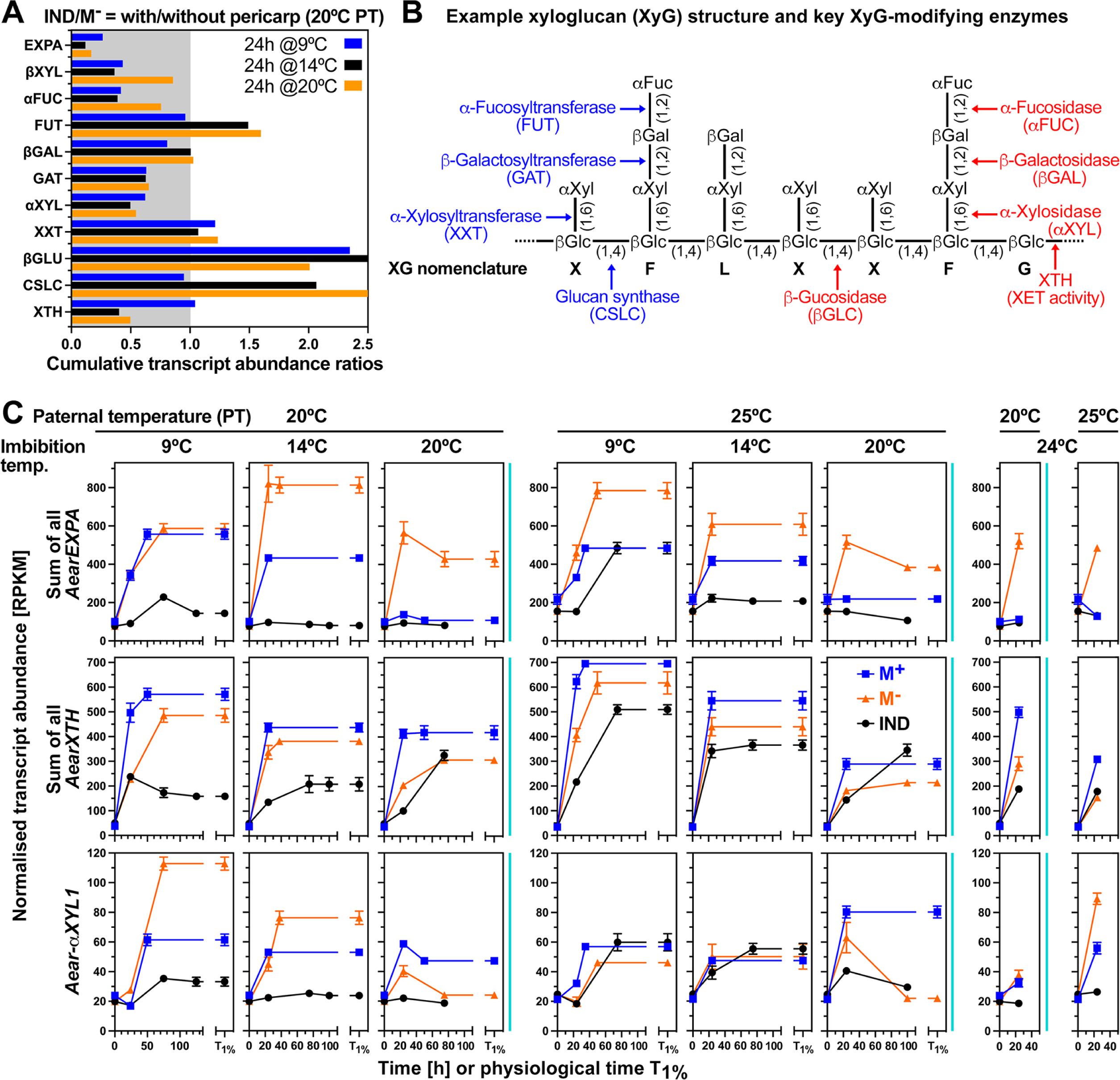
Transcript abundance patterns (RNA-seq) of *Aethionema arabicum* cell-wall remodeling protein genes in seeds of imbibed dimorphic diaspores (M^+^ seeds, IND fruits) and bare M^−^ seeds. A, Effect of the pericarp on the expression ratios of expansin A and xyloglucan-related cell-wall remodeling protein genes in the M^−^ seeds of 24 h imbibed IND fruits and isolated M^−^ seeds. B, Xyloglucan remodeling is achieved by a battery of enzymes specifically targeting different bonds of xyloglucan structure as indicated. Among them are xyloglucan *endo*-transglycoylases/hydrolases (XTHs) with xyloglucan *endo*-transglycoylase (XET) enzyme activity (Holloway et al., 2021). C, Transcript abundance patterns of *Ae. arabicum* expansins A, XTHs and the α-xylosidase *Aear-αXYL1* in M^+^ seeds, IND fruits and isolated M^−^ seeds from two parental temperature regimes (20°C versus 25°C) at four different imbibition temperatures (9, 14, 20 and 24°C). For other expansin and xyloglucan-related genes and *Ae. arabicum* gene IDs see Supplemental Figure S16 or the Expression Atlas (https://plantcode.cup.uni-freiburg.de/aetar_db/index.php); for RNAseq single values see the Expression Atlas or Supplemental Data Set 1. WGCNA modules (Figure 3) are indicated by the vertical color lines next to the graphs. Mean ± SEM values of 3 replicates each with 60-80 seeds are presented.

In contrast to the observed hypoxia-induced expression of many genes, hypoxia inhibited the induction of the expansin, XTH and *αXYL1* cell wall-remodeling gene expression in imbibed bare M^−^ seeds (Figure 8A). Compared to hypoxia, ABA was far less effective and did not appreciably inhibit the induction of the accumulation of *AearEXPA2*, *AearEXPA9*, *AearXTH4*, *AearXTH16a* and *AearαXYL1* transcripts. The pericarp effect on the gene expression patterns (IND fruits versus bare M^−^ seeds) observed in the transcriptome analysis was confirmed in the completely independent RT-qPCR experiment for these cell wall-remodeling genes and for most of the total of 32 genes investigated (Supplemental Figure S13C). While for the representative genes for each WGCNA module pericarp extract did not affect the gene expression of bare M^−^ seeds (Supplemental Figure S13C), hypoxia conditions affected it for about half of the representative genes (Figure 8B). Correlation analysis between the pericarp effect imposed to M^−^ seeds in imbibed IND fruits and the hypoxia and ABA effects on imbibed bare M^−^ seeds was conducted for the 32 genes of the RT-qPCR experiment (Figure 8C). This revealed strong linear relationships (R^2^ 0.7-0.8) for hypoxia versus pericarp, and hypoxia+ABA versus pericarp, but not for ABA alone versus pericarp. Taken together, hypoxia generated by the pericarp seems to be the most important mechanism of the pericarp-mediated dormancy of IND fruits, it seems to act upstream of ABA and affects the gene expression in M^−^ seed encased by the pericarp to control the observed distinct dormancy and germination responses at the different imbibition temperatures.

## Discussion

### *Aethionema arabicum* seed and fruit dimorphism: large-scale molecular data sets reveal diaspore bet-hedging strategy mechanisms in variable environments

Ambient temperature during seed reproduction (parental temperature) and after dispersal (including imbibition temperature) is a major determinant for fecundity, yield, seed germinability (i.e. nondormancy versus dormancy of different depth), and environmental adaptation. The effect of temperature variability has been well-studied in monomorphic annual plants and mechanisms underpinning the germinability of the dispersed seeds were elucidated (Donohue et al., 2010; Finch-Savage and Footitt, 2017; Fernandez-Pascual et al., 2019; Iwasaki et al., 2022; Zhang et al., 2022). In contrast to monomorphic species, very little is known about the morphological and molecular mechanisms that facilitate survival of heteromorphic annual species. Diaspore heteromorphism is the prime example of bet-hedging in angiosperms and is phenologically the production by an individual plant of two (dimorphism) or more seed/fruit morphs that differ in morphology, germinability, and stress ecophysiology to ‘hedge its bets’ in variable (unpredictable) environments (Imbert, 2002; Baskin et al., 2014; Gianella et al., 2021).

An advantage of *Ae. arabicum* as a model system is that it exhibits true seed and fruit dimorphism with no intermediate morphs (Lenser et al., 2016). Our earlier work revealed molecular and morphological mechanisms underlying the dimorphic fruit and seed development (Lenser et al., 2018; Wilhelmsson et al., 2019; Arshad et al., 2021), the distinct dispersal properties of the M^+^ seed and IND fruit morphs (Arshad et al., 2019; Arshad et al., 2020; Nichols et al., 2020), and the adaptation to specific environmental conditions (Mohammadin et al., 2017; Bhattacharya et al., 2019; Merai et al., 2019; Merai et al., 2023). Transcriptome and imaging analyses of the seed coat developmental program of the mucilaginous *Ae. arabicum* M^+^ seed morph revealed that it resembles the ‘default’ program known from the mucilaginous seeds of *A. thaliana*, *Lepidium sativum*, and other Brassicaceae (Graeber et al., 2014; Scheler et al., 2015; Lenser et al., 2016; Arshad et al., 2021; Steinbrecher and Leubner-Metzger, 2022). In contrast to this, the non-mucilaginous *Ae. arabicum* M^−^ seed morph resembles *A. thaliana* seed mucilage mutants and thereby highlights that the dimorphic diaspores enable the comparative analysis of distinct developmental programs without the need for mutants. *Arabidopsis thaliana* accessions differ in depth of their primary seed dormancy, reaching from the deep physiological to shallow dormancy (Cadman et al., 2006; Barrero et al., 2010; Finch-Savage and Footitt, 2017); and *L. sativum* seeds are completely non-dormant (Graeber et al., 2014). During seed imbibition, distinct transcriptional and hormonal regulation either leads to the completion of germination or to dormancy maintenance for which ABA metabolism and signaling, *DOG1* expression, downstream cell wall remodeling and seed coat properties are key components (Finch-Savage and Leubner-Metzger, 2006; Graeber et al., 2014; Footitt et al., 2020; Iwasaki et al., 2022). The transcriptome and hormone data for *Ae. arabicum* M^+^ seeds confirmed these mechanisms and their dependence on the imbibition temperature to either mount a germination or a dormancy program typical for mucilaginous seeds (Cadman et al., 2006; Scheler et al., 2015; Iwasaki et al., 2022). The *Ae. arabicum* dimorphic diaspore comparison of these M^+^ seeds to IND fruits (and the bare M^−^ seeds) revealed how the pericarp of the IND fruit morph imposes the observed coat dormancy.

### Pericarp-imposed dormancy: comparative analyses of indehiscent fruit and seed morph germinability reveals mechanisms and roles in thermal responses

The typical Brassicaceae fruit is dehiscent, opens during fruit maturation (dehiscence, *A. thaliana* seed, *Ae. arabicum* M^+^ seed morph), and is considered to represent the ancestral fruit type (Mühlhausen et al., 2013). Nevertheless, monomorphic species that disperse various indehiscent fruit types by abscission evolved many times independently within the Brassicaceae. Different roles of the pericarp in these dry indehiscent Brassicaceae diaspores (siliques and silicles) were identified, including dispersal by wind, persistence in the seed bank, retaining seed viability, delaying water uptake, releasing allelochemicals, and imposing coat dormancy (Mamut et al., 2014; Lu et al., 2015b; Lu et al., 2017b, a; Sperber et al., 2017; Mohammed et al., 2019; Khadka et al., 2020). Many of these monomorphic Brassicaceae species with indehiscent fruits are desert annuals. Their indehiscent diaspores were not investigated for the molecular mechanisms responding to distinct parental and imbibitional temperatures. The *Ae. arabicum* dimorphic diaspore system with its single-seeded IND fruit morph, provides an excellent system for investigating these mechanisms.

The pericarp of mature *Ae. arabicum* IND fruits is dead tissue that contains high amounts of ABA, OPDA, JA, JA-Ile, and SA, as well as degradation products of ABA and IAA. Leaching of these and other compounds into the fruit’s proximal environment could have roles in allelopathic interactions, as described for the dead pericarp of other species (Grafi, 2020; Khadka et al., 2020). Leaching of pericarp inhibitors into the encased seed could also delay fruit germination or confer ‘chemical coat dormancy’, as it was demonstrated for pericarp-derived ABA in *Lepidium draba* (Mohammed et al., 2019), *Beta vulgaris* (Ignatz et al., 2019), and *Salsola komarovii* (Takeno and Yamaguchi, 1991). Pericarp extracts of *Ae. arabicum* IND fruits as well as ABA delayed M^−^ seed germination, but they could not fully mimic the pericarp-imposed dormancy and effect on gene expression (Figure 5). Acting as a mechanical restraint to water uptake and/or radicle protrusion is another way by which the pericarp may delay germination or confer ‘mechanical coat dormancy’ (Sperber et al., 2017; Steinbrecher and Leubner-Metzger, 2017). The *Ae. arabicum* IND pericarp is water-permeable (Lenser et al., 2016), and we showed here that it weakens during imbibition. *Aethionema arabicum* pericarp-imposed dormancy was enhanced by the lower parental temperature (20°C, 20IND fruits) as compared to the higher parental temperature (25°C, 25IND fruits). This altered the pericarp biochemically and its mechanical resistance, which was higher in 20IND pericarps (Figure 6), as was the pericarp-imposed dormancy (Figure 1).

The role of the pericarp and other seed-covering structures in limiting oxygen availability to the embryo (hypoxia) is another mechanism for coat-imposed dormancy (Benech-Arnold et al., 2006; Mendiondo et al., 2010; Dominguez et al., 2019). Comparative transcriptome and hormone analyses (IND, M^−^, M^+^) identified that upregulated expression of hypoxia-responsive genes is a hallmark of imbibed IND fruits (i.e., in M^−^ seeds encased by the dead pericarp) as compared to M^+^ seeds and bare M^−^ seeds (Figure 7). Identified hypoxia-responsive genes include the hypoxia-induced ERF-VII TF *AearERF71/73* and the fermentation genes *AearADH1a* and *AearPDC2*, but not *AearADH1b* and *AearPDC1*. While the *AearADH1a*, *AearPDC2,* and *AtPDC1* gene 5’-regulatory region contain ERF73 and HRPE motifs, they do not contain G-box/ABRE motifs as this is the case for the *AearADH1b*, *AearPDC1,* and *AtADH1* genes. The relative importance of *AtERF71/73*, ABA-related, and other TFs in the hypoxia-induced expression of the *AtADH1* gene in *A. thaliana* is not completely resolved (Lu et al., 1996; Kürsteiner et al., 2003; Gomez-Porras et al., 2007; Yang et al., 2011; Papdi et al., 2015; Gasch et al., 2016; Seok et al., 2022). For *Ae. arabicum,* we speculate that the observed duplication of the ADH genes, the differences in the *cis*-regulatory motifs, the accumulation of *AearADH1a* and *AearPDC2* transcripts in M^−^ seeds inside IND fruits, and pericarp-mediated hypoxia leading to PDC-ADH catalyzed ethanolic fermentation constitute a morph-specific adaptation that contributes to the increased dormancy of IND fruits.

### Morphological and hormonal regulation: pericarp-ABA interactions as a key mechanism for distinct post-dispersal dimorphic diaspore responses to environmental cues

Earlier work with dormant barley grains and sunflower demonstrated that hypoxia, imposed either artificially or by the maternal seed covering structures (barley glumellae, sunflower pericarp), interfered with ABA metabolism and increased embryo ABA sensitivity (Benech-Arnold et al., 2006; Mendiondo et al., 2010; Andrade et al., 2015; Dominguez et al., 2019). In barley, this included transient ABA accumulation and ABI5 gene expression during dormancy maintenance. In sunflower, the pericarp-imposed dormancy was associated with increased embryo sensitivity to hypoxia and ABA, but with no change in embryo ABA content (Dominguez et al., 2019). As in *Ae. arabicum* pericarp (Figure 6), sunflower pericarp also contained considerable amounts of ABA, SA, OPDA, JA, and JA-Ile (Andrade et al., 2015). In general, their contents declined during imbibition in both species, except ABA, which accumulated transiently in the sunflower pericarp, but declined in the dead pericarp of *Ae. arabicum*. The *Ae. arabicum* IND fruit versus M^−^ seed comparison revealed the decisive role of the pericarp and ABA in narrowing the germination-permissive window (Figure 1). Using the three-way transcriptome and hormone comparison (IND, M^−^, M^+^), we could identify mechanisms not existing in monomorphic species. These include a very clear temperature-dependent up-regulation of ABA biosynthesis genes (including *AearNCED6*) and down-regulation of ABA 8’-hydroxylase (including *AearCYP707A3*) in M^−^ seeds within imbibed IND fruit morphs as compared to imbibed bare M^−^ seeds and the M^+^ seed morphs (Figure 5). At the 9°C and 14°C imbibition temperatures, the resultant ABA accumulation in M^−^ seeds inside IND fruits was especially elevated in the more dormant 20IND fruits as compared to the less dormant 25IND fruits.

In agreement with a pericarp-enhanced ABA sensitivity of M^−^ embryos inside IND fruits, pericarp-enhanced expression of numerous ABA-related TFs including *AearAREB3*, *AearABI5*, *AearABF1*, and *AearGBF3* (Figure 8) became evident during imbibition. Further, the presence of the pericarp affected the expression patterns for major ABA signaling genes, including for ABA receptors, PP2Cs, and SnRK2, which control the ABA-related TFs. The control of germinability by ABA signaling is, in part, achieved by regulating the expression of downstream cell wall remodeling genes (e.g. Finch-Savage and Leubner-Metzger, 2006; Barrero et al., 2010; Shigeyama et al., 2016; Steinbrecher and Leubner-Metzger, 2017; Holloway et al., 2021; Steinbrecher and Leubner-Metzger, 2022). Their expression in imbibed *Ae. arabicum* IND fruits, M^+^ and M^−^ seeds also exhibited pericarp– and temperature-dependent patterns (turquoise module in most cases), which is mainly mediated by hypoxia affecting ABA sensitivity and gene expression (Figures 8 and 10).

The presented new, comprehensive molecular datasets on responses of dispersed dimorphic diaspores to ambient temperature, together with previous work on fruit/seed development (Lenser et al., 2018; Wilhelmsson et al., 2019; Arshad et al., 2021), highlights *Ae. arabicum* as the best experimental model system for heteromorphism so far. It provides a growing potential to understand developmental control and plasticity of fruit and seed dimorphism and it’s underpinning molecular, evolutionary, and ecological mechanisms as adaptation to environmental change. The comparative analysis of the M^+^ seed morph, the IND fruit morph, and the bare M^−^ seed revealed morphological, hormonal, and gene regulatory mechanisms of the pericarp-imposed dormancy. The dimorphic diaspores integrate parental and imbibition temperature differently, involving distinct transcriptional changes and ABA-related regulation. The *Ae. arabicum* web portal (https://plantcode.cup.uni-freiburg.de/aetar_db/index.php) with its genome database and gene expression atlas comprises published transcriptome results (this work and Merai et al., 2019; Wilhelmsson et al., 2019; Arshad et al., 2021), is open for dataset additions, makes the data widely accessible, and providing a valuable source for future work on diaspore heteromorphism.

## Materials and Methods

### Plant material, experimental growth conditions, and germination assays

Plants of *Aethionema arabicum* (L.) Andrz. ex DC. were grown from accessions TUR ES1020 (from Turkey) (Mohammadin et al., 2017; Mohammadin et al., 2018; Merai et al., 2019), in Levington compost with added horticultural grade sand (F2+S), under long-day conditions (16 h light/ 20°C and 8 h dark/ 18°C) in a glasshouse. Upon onset of flowering, plants were transferred to distinct parental temperature regimes (20°C versus 25°C) during reproduction in otherwise identical growth chambers as described (Supplemental Figure S1A). Mature M^+^ seeds and IND fruits were harvested (Supplemental Table S1), further dried over silica gel for a week and either used immediately or stored at –20°C in air-tight containers. For germination assays, dry mature seeds (M^+^ or M^−^) or IND fruits were placed in 3 cm Petri dishes containing two layers of filter paper, 3 ml distilled water and 0.1% Plant Preservative Mixture (Plant Cell Technology). Temperature response profiles (Figure 1C) were obtained by incubating plates on a GRD1-LH temperature gradient plate device (Grant Instruments Ltd., Cambridge, UK). Subsequent germination assays were conducted by incubating plates in MLR-350 Versatile Environmental Test Chambers (Sanyo-Panasonic) at the indicated imbibition temperature and 100 µmol • s^−1^ • m^−2^ continuous white light (Lenser et al., 2016). Germination assays under hypoxia conditions compressed air (UN1002, BOC Ltd., Woking, UK) and oxygen-free nitrogen (BOC UN1066) were mixed to generate a 4.5±0.2% oxygen atmosphere in hypoxia chambers (Stemcell Technolgies, Waterbeach, Cambridge, UK) with the plates (14°C, continuous white light). Seed germination, scored as radicle emergence, of 3 biological replicates each with 20 to 25 seeds or fruits were analyzed.

### Multispectral Imaging (MSI), biomechanical and pericarp extract assays

Multispectral imaging was performed with a VideometerLab (Mark4, Series 11, Videometer A/S, Denmark). Images were transformed using normalized canonical discriminant analysis (nCDA) to compare the two parental temperatures. Biomechanical properties of the fruit coats were measured using a universal material testing machine (ZwickiLine Z0.5, Zwick Roell, Germany). Fruits were imbibed for 1 h before cutting them in half (fruit half covering the micropylar end of the seed and non-micropylar end of the seed) and re-dried overnight. Seeds were removed from the pericarps, and a metal probe with a diameter of 0.3 mm was driven into the sample at a speed of 2 mm/min while recording force and displacement. Tissue resistance was determined to be the maximal force from the force-displacement curve. Pericarp extract (PE) was obtained as aqueous leachate by incubating ca. 0.5 g IND pericarp in 15 ml H_2_O on a shaker for 2 hours, followed by cleaning it using a 0.2 µm filter. Germination assays were conducted by comparing PE, H_2_O (control), *cis,trans*-S(+)-ABA (ABA; Duchefa Biochemie, Haarlem, The Netherlands), salicylic acid (SA; Alfa Aesar, Lancashire, UK), *cis*-(+)-12-oxophytodienoic acid (OPDA; Cayman Chemical, MI), (-)-jasmonic acid (JA; Cayman Chemical, MI, USA), or its isoleucine conjugate (JA-Ile; Cayman Chemical) at the concentrations indicated.

### RNA-seq and quantitative reverse transcription-PCR (RT-qPCR)

Sampling of dry or imbibed M^+^ seeds, M^−^ seeds, and IND fruits for molecular analyses was as described in the sampling scheme (Supplemental Figure S1). Biological replicates of samples each corresponding to 20 mg dry weight of seed material, were pulverized in liquid N_2_ using mortar and pestle. Extraction of total RNA was performed as described by Graeber et al. (2011). RNA quantity and purity were determined using a NanoDrop™ spectrophotometer (ND-1000, ThermoScientific™, Delaware, USA) and an Agilent 2100 Bioanalyzer with the RNA 6000 Nano Kit (Agilent Technologies, CA, USA) using the 2100 Expert Software to calculate RNA Integrity Number (RIN) values. Four (RT-qPCR) or three (RNAseq) biological replicates of RNA samples were used for downstream applications (Sample naming scheme: Supplemental Table S2). Sequencing was performed at the Vienna BioCenter Core Facilities (VBCF) Next Generation Sequencing Unit, Vienna, Austria (www.vbcf.ac.at). RNA-seq libraries were sequenced in 50 bp single-end mode on Illumina® HiSeq 2000 Analyzers using the manufacturer’s standard module generation and sequencing protocols. The overall sequencing and mapping statistics for each library and the read counts are presented in Supplemental Data Set 1. RNA for RT-qPCR was extracted in an independent experiment using the RNAqueous Total RNA Isolation Kit with the addition of the Plant RNA Isolation Aid (Ambion, Thermo Fisher Scientific, Basingstoke, UK), followed by treatment with DNaseI (QIAGEN Ltd., Manchester, UK) and precipitation in 2 M LiCl. Precipitated RNA was washed in 70% ethanol and resuspended in RNase-free water. RT-qPCR was conducted and analyzed as described (Graeber et al., 2011; Untergasser et al., 2012; Arshad et al., 2021) using primer sequences and reference genes listed in Supplemental Table S3.

### Analyses of transcriptome data

Transcriptome assembly, data trimming, filtering, read mapping and feature counting, and DEG detection were performed as previously described (Wilhelmsson et al., 2019; Arshad et al., 2021). Principal component analysis (PCA) was performed using the built-in R package “prcomp” (www.r-project.org) on log(x+1) transformed RPKM values for 22200 genes with non-zero values in at least one sample. Sample replicate RPKM values were averaged for 45 treatments and WGCNA (Zhang and Horvath, 2005) implementation (Langfelder and Horvath, 2008) in R was performed on log_2_(x+1) transformed RPKM values for 11260 genes whose average expression was >4 RPKM across all samples. The function blockwiseModules was used with default settings, other than to create a signed hybrid network distinguishing between positive and negative Pearson correlations using a soft power threshold of 24, minModuleSize of 50, mergeCutHeight of 0.25, and pamRespectsDendro set to False in single block. Module membership and significance for each gene were calculated (Pearson correlation with module eigengene) (Supplemental Data Set 2). PCA analysis (Wickham, 2016) for the 11260 genes was performed as outlined above with transposed data. Module eigengene expression was correlated with sample traits using Pearson correlation. GO term enrichment in module gene lists was calculated using the R package topGO (Alexa and Rahnenfuhrer, 2023) using the elim or classic method with Fisher’s exact test. Geneious 8.1.9 (https://www.geneious.com) was used to visualize motif positions. Gene identifier and symbols (Supplemental Table S2) are according to earlier publications of the *Ae. arabicum* genome and transcriptome (Haudry et al., 2013; Merai et al., 2019; Nguyen et al., 2019; Wilhelmsson et al., 2019; Arshad et al., 2021) and the *Ae. arabicum* web portal (https://plantcode.cup.uni-freiburg.de/aetar_db/index.php) links this to the current (Fernandez-Pozo et al., 2021) and future genome DB and gene expression atlas.

### Gene promoter analyses

Promoter motif enrichment in the start codon –1000 to +100 bp region was analyzed using the Analysis of Motif Enrichment tool (McLeay and Bailey, 2010) using MEME Suite (https://meme-suite.org/) (Bailey et al., 2015) to identify enrichment of motifs from the ArabidopsisDAPv1 database (O’Malley et al., 2016). Input sequences (module gene list) were compared to control sequences (all promoter sequences) using average odds score, Fisher’s exact test, fractional score threshold of 0.25, E-value cutoff of 10, and 0-order background model. FIMO (Grant et al., 2011) on MEME Suite was used to scan sequences for chosen motifs. Chord diagram was drawn using R package “circlize” (Gu et al., 2014).

### Phytohormone quantification

For quantification of jasmonates (JA, JA-Ile and *cis*-OPDA), auxins (IAA and its catabolite oxIAA), abscisates (ABA, PA and DPA) and salicylic acid (SA), internal standards, containing 20 pmol of [^2^H_4_]SA and [^2^H_5_]OPDA, 10 pmol each of [^2^H_6_]ABA, [^2^H_6_]JA and [^2^H_2_]JA-Ile, and 5 pmol each of [^2^H_3_]PA, [^2^H_3_]DPA, [^13^C_6_]IAA and [^13^C_6_]oxIAA (all from Olchemim Ltd, Czech Republic), and 1 ml of ice-cold methanol: water (10:90, v/v) were added to 10 mg of freeze-dried and homogenized samples. Sample mixtures were homogenized using an MM400 vibration mill for 5 min at 27 Hz (Retsch Technology GmbH, Germany), sonicated for 3 min at 4 °C using an ultrasonic bath, and then extracted for 30 min (15 rpm) at 4°C using a rotary disk shaker. Samples were centrifuged at 20,000 rpm (15 min, 4 °C), the supernatant purified using pre-equilibrated Oasis HLB cartridges (1 cc, 30 mg, Waters), and evaporated to dryness under nitrogen (30°C) (Flokova et al., 2014). The evaporated samples were reconstituted in 40 μl of the mobile phase (15% acetonitrile, v/v) and analyzed by UHPLC-ESI-MS/MS as described by Šimura et al. (2018). All phytohormones were detected using a multiple-reaction monitoring mode of the transition of the precursor ion to the appropriate product ion. Masslynx 4.1 software (Waters, Milford, MA, USA) was used to analyze the data, and the standard isotope dilution method (Rittenberg and Foster, 1940) was used to quantify the phytohormone levels. Five independent biological replicates were performed.

## Statistical analysis

Germination data were evaluated by comparing final germination percentage (G_max_) and germination rate (speed). Germination curve fits and T_50%_ were calculated with GERMINATOR (Joosen et al., 2010). An unpaired t-test was used to compare the mean values for tissue resistance (biomechanical analysis of pericarp) of the two parental temperatures. All statistical analyses were performed in GraphPad Prism (v. 8.01, GraphPad Software Inc., San Diego, California, USA).

## Data availability and accession numbers

The RNAseq data discussed in this publication have been deposited at the NCBI Sequencing Read Archive (SRA), BioProjects PRJNA611900 (dry seed) and PRJNA639669 (imbibed seed), accessible at https://www.ncbi.nlm.nih.gov/sra; metadata about the samples are also available as part of this publication (Supplemental Data Set 1). Further, normalized transcriptome data from this study and associated previous studies (Merai et al., 2019; Wilhelmsson et al., 2019; Arshad et al., 2021) can be accessed and visualized at the *Ae. arabicum* web portal (https://plantcode.cup.uni-freiburg.de/aetar_db/index.php). For *Ae. arabicum* gene IDs see Supplemental Figures S15, S16, Supplemenal Table 2 or the Expression Atlas (https://plantcode.cup.uni-freiburg.de/aetar_db/index.php); for RNAseq single values see the Expression Atlas or Supplemental Data Set 1. All other data presented or analyzed in this published article are available online through the supplements.

## Supplemental data

The following materials are available in the online version of this article.

**Supplemental Figure S1.** Large-scale diaspore production experiment and sampling

**Supplemental Figure S2.** *Aethionema arabicum* gene expression atlas

**Supplemental Figure S3.** PCA of transcriptome data (RNA-seq)

**Supplemental Figure S4.** Expression of WGCNA modules

**Supplemental Figure S5.** Temperature responses and ABA metabolism

**Supplemental Figure S6.** Hormone contents in response to temperatures

**Supplemental Figure S7.** Effects of pericarp extract on germination and gene expression

**Supplemental Figure S8.** Pericarp biomechanics

**Supplemental Figure S9.** Transcription factors and promoter motifs of fermentation genes

**Supplemental Figure S10.** Detail comparison of fermentation gene promoters

**Supplemental Figure S11.** Transcript abundance patterns of the seed-specific Perl’s pathway

**Supplemental Figure S12.** Transcript abundance patterns of hypoxia-regulated genes

**Supplemental Figure S13.** RT-qPCR analysis of hypoxia and ABA effects

**Supplemental Figure S14.** Transcript abundance patterns of ABA-related genes

**Supplemental Figure S15.** Transcript abundance patterns of transcription/translation genes

**Supplemental Figure S16.** Transcript abundance patterns of cell wall remodeling genes

**Supplemental Table S1.** *Aethionema arabicum* seed and fruit harvest results

**Supplemental Table S2.** *Aethionema arabicum* gene IDs, symbols and modules

**Supplemental Table S3.** RT-qPCR primer used

**Supplemental Data Set 1.** Gene annotations, DEGs, RPKM values, raw counts, overall sequencing and mapping statistics for each library, and SRA accessions

**Supplemental Data Set 2.** WGCNA modules and GO term enrichment analysis

**Supplemental Data Set 3.** Promoter enrichment analysis (AME) and chord diagram data

**Supplemental Data Set 4.** Promoter motif scanning (FIMO)

## Acknowledgements

We acknowledge the support of Katja Sperber, Teresa Lenser, Samik Bhattacharya, Sara Mayland-Quellhorst, Setareh Mohammadin, Safina Khan, Christopher Grosche, Giles Grainge, Thomas Holloway, Kazumi Nakabayashi and Michael Ignatz in discussions, harvesting of diaspores and other tasks. The authors are grateful for the excellent support by the Next Generation Sequencing Unit of the Vienna BioCenter Core Facilities (VBCF). We acknowledge the support of DataPLANT (NFDI 7/1 – 42077441) as part of the German National Research Data Infrastructure.

## Author Contribution Statement

K.G., K.M., G.T., M.S., O.M.S., M.E.S., O.N., S.A.R. and G.L.-M. conceived the project and conceptualized the work. J.O.C., P.K.I.W. and K.G. performed the majority of experiments; K.G., Z.M. and J.O.C. prepared and handled samples; J.O.C., P.K.I.W., K.K.U., W.A., S.A.R. and G.L.-M. performed the transcriptomics data analysis. J.O.C., T.S. and M.P. performed RT-qPCR and hypoxia analyses. J.O.C., T.S. and M.P. performed germination, biomechanical experiments, and multispectral imaging analyses. N.F.-P., J.O.C. and S.A.R. developed and implemented the gene expression atlas. T.-P.N., K.G., W.A., K.M. and M.E.S. conducted the environmental simulation experiment and generated the plant material. I.P., D.T., O.N. and M.S. performed the hormone analysis. J.O.C. and G.L.-M. wrote the manuscript. All authors read and commented on the manuscript.

